# Epigenetic machinery is functionally conserved in cephalopods

**DOI:** 10.1101/2021.11.18.469068

**Authors:** Filippo Macchi, Eric Edsinger, Kirsten C. Sadler

## Abstract

Epigenetic regulatory mechanisms are divergent across the animal kingdom, yet little is known about the epigenome in non-model organisms. Unique features of cephalopods make them attractive for investigating behavioral, sensory, developmental and regenerative processes, but using molecular approaches in such studies is hindered by the lack of knowledge about genome organization and gene regulation in these animals. We combined bioinformatic and molecular analysis of *Octopus bimaculoides* to identify gene expression signatures for 12 adult tissues and a hatchling, and investigate the presence and pattern of DNA methylation and histone methylation marks across tissues. This revealed a dynamic gene expression profile encoding several epigenetic regulators, including DNA methylation maintenance factors that were highly conserved and functional in cephalopods, as shown by detection of 5-methyl-cytosine in multiple tissues of octopus, squid and bobtail squid. WGBS of octopus brain and RRBS from a hatchling revealed that less than 10% of CpGs are methylated, highlighting a non-random distribution in the genome of all tissues, with enrichment in the bodies of a subset of 14,000 genes and absence from transposons. Each DNA methylation pattern encompassed genes with distinct functions and, strikingly, many of these genes showed similar expression levels across tissues. In contrast to the static pattern of DNA methylation, the histone marks H3K27me3, H3K9me3 and H3K4me3 were detected at different levels in diverse cephalopod tissues. This suggests the methylome and histone code cooperate to regulate tissue specific gene expression in a way that may be unique to cephalopods.

## Introduction

Epigenetic modifications to histones and DNA are conserved features that regulate tissue-specific profiles of gene expression and repress transposable elements (TEs) [1]. There is great diversity across the animal kingdom in how the epigenome accomplishes these complex and important functions, yet mechanistic studies on epigenetic patterning and function have largely focused on a few model organisms [2]. While this has been extremely fruitful in elucidating mechanisms of epigenome patterning, regulation and function, they do not provide a comprehensive understanding of how epigenetic marks are patterned or how they regulate genes, TEs or genome structure across the evolutionary tree.

Examining DNA methylation in diverse animal species provides an illustrative example of how incorporating organismal diversity into epigenetic studies expands the epigenetic lexicon. In vertebrates, the vast majority of CpGs are methylated, with an asymmetric distribution throughout the genome, including enrichment in intergenic regions, exclusion from CpG rich promoters, and a varied pattern on gene bodies [3]. Heavy decoration of repetitive sequences with 5-methyl cytosine (5mC) reflects its important function in suppressing the expression of CpG dense transposons [4]. In contrast, the initial studies on canonical invertebrate model organisms - *Caenorhabditis elegans* and *Drosophila melanogaster* – showed there was a complete lack of DNA methylation [5-7]. The result of this is that detailed functional analysis of DNA methylation in animals has been largely carried in vertebrate model organisms and human samples, creating a large gap in understanding of how the DNA methylome is patterned or what its function is in other animals.

The advance of sequencing technologies has massively expanded the diversity of organisms with fully sequenced genomes, allowing examination of DNA methylation patterns in animals from across the evolutionary tree [3, 8-11]. This revealed that most invertebrates have either no or low levels of CpG methylation. Moreover, the pattern of 5mC distribution is very different in animals with low levels of methylation compared to vertebrate genomes, with methylated CpGs enriched in gene bodies and heavily methylated genes tend to be expressed at higher levels than unmethylated genes [3, 8, 9, 12-16]. Exceptions to these patterns abound; for example, the methylome in a sponge is highly similar to vertebrates [17, 18], the sea squirt *Ciona intestinalis* shows a high level of gene body methylation with an intermediate methylation pattern on repeats [9, 10, 19] and there are dynamic changes in the DNA methylation pattern during oyster development [15, 20], compared to the largely static pattern across tissues and developmental stages in vertebrates.

Unique features of the *Octopus bimaculoides* methylome was highlighted by a recent study using whole genome bisulfite sequencing (WGBS) to analyze brain methylation patterns in diverse animal species [16]. In most animals, TE abundance correlates with genome size, and since DNA methylation serves to keep transposons in check in vertebrates and plants [12], it has been hypothesized that DNA methylation is a key to suppressing TE expression in organisms with large genomes and high transposon burden [21]. This is not the case in *O. bimaculoides*, where half the genome is populated by repetitive elements similarly to vertebrates [22]. Interestingly, the level of DNA methylation is not proportional to the transposon load in this genome [16]. This raises the important question of whether the DNA methylation pattern observed in octopus is conserved in other cephalopods and warrants investigation of the function of DNA methylation in these animals.

In species which have DNA methylation, the pattern of CpG methylation needs to be copied to the daughter strand during DNA replication. This is carried out by the DNA methyltransferase DNMT1 and by UHRF1, which recognizes hemi-methylated DNA generated during DNA replication and recruits DNMT1 [23-28]. UHRF1 also functions as a reader of the histone code, including the canonical heterochromatin mark, trimethylated histone H3 lysine 9 (H3K9me3) [29-34]. The DNMT1-UHRF1-complex represents the core DNA methylation machinery and contributes to establishing heterochromatin domains. Recent studies reported high conservation of DNMTs, UHRF1 and TET proteins across diverse phylogenetic groups of animals including annelid worms [18], sponges [17, 18], mollusks [14, 35], and other invertebrates [3, 16]. This indicates that although 5mC is not ubiquitous throughout the animal kingdom, in those cases where it is present, it plays diverse, albeit incompletely understood, roles.

Cephalopods represent an emerging model system with multiple studies utilizing squid, bobtail squid and octopus for uncovering novel mechanisms of RNA editing, highly complex behavioral regulation, remarkable regenerative capacity, and genome evolution [36-44]. Recent advances in embryo cultivation, standardized aquaculture protocols [45], and genome editing [46] have advanced the utility of cephalopods as new model organisms [47]. Transcriptomic profiling of mollusk embryos [48], brain [49, 50] and multiple tissues of *O. bimaculoides* adults [22] have revealed that key genes having a role in development and neurological processes are highly conserved in these animals. However, a comprehensive analysis of tissue-specific gene expression profiles in cephalopods has not been reported. Moreover, despite significant strides in cephalopod research and few studies reporting the presence of DNA methylation in some species of octopus [16, 51, 52], there is virtually nothing known about the epigenetic marks that contribute to the regulation of tissue-specific gene expression profiles, transposon suppression or genome organization in cephalopods.

We address this using transcriptomic and methylome datasets and biochemical approaches to analyze *O. bimaculoides* tissue specific gene expression, methylation patterning and histone modifications. We show that DNA methylation and histone modifications are present in multiple tissues of three cephalopod species. Methylome analysis in octopus shows that while methylated CpGs account for less than 10% of all CpGs in the genome and are virtually absent from repetitive DNA and transposons. Methylated CpGs are clustered on the gene bodies of a distinct set of genes which are highly expressed across tissues. This shows that CpG methylation and histone methylation are prominent features of the cephalopod epigenome and suggests that the pattern of DNA methylation is set by characteristics of the genome that are maintained across cell types.

## Materials and Methods

### Animal husbandry and sample collection

Animals were maintained in a circulating natural sea water aquaculture facility at the Marine Resources Center at the Marine Biological Laboratory and all experiments were performed according to the current policy for the use of cephalopods at Marine Biology Laboratories (MBL, https://www.mbl.edu/policies/files/2018/07/J1.10-CEPH-Policies-Procedures-Jan-1-31-2020-fillable-form.pdf). In brief, adult male octopus *O. bimaculoides*, adult male bobtail squid *Euprymna berryi* and adult squid *Doryteuthis pealeii* (unknown sex) were anesthetized in 3% ethanol in natural sea water for 10 minutes and tissues were dissected, flash frozen in liquid nitrogen, and stored at -80°C. One whole *O. bimaculoides* hatchling at developmental stage of 30 days post-fertilization (30 dpf hatchling) was anesthetized as above and flash frozen.

### RNA and DNA extraction

Frozen tissues were ground using a mortar and pestle cooled with liquid nitrogen and placed in dry ice. 15 mg of tissue powder was used to extract either RNA or DNA. RNA was extracted using Trizol (Invitrogen) following the manufacturer’s instructions with some modifications. Briefly, during precipitation in isopropanol, 10 µg of Glycoblue (Thermo Fisher Scientific) was added and precipitation was performed overnight at -20°C followed by 1 hour centrifuge at 12000g at 4°C. RNA was resuspended in water and used in the following procedures. Genomic DNA (gDNA) was extracted by using a DNA extraction buffer (10 mM Tris-HCl pH9, 10 mM EDTA, 200 mM NaCl, 0.5% DSD, 200 µg/ml proteinase K) and over-night incubation at 65°C, followed by RNAase treatment with 2 mg/ml PureLink™ RNase A (Invitrogen) for 2 h at 37°C. Then, 0.25 v/v of 5 M potassium acetate (CH_3_CO_2_K) was added and the sample centrifuged at 12,000 x g at room temperature to precipitate proteins. 1:1 v/v of isopropanol was added to the supernatant and incubated at -20°C overnight and DNA precipitated by centrifuge at 12,000 x g at room temperature for 15 minutes. DNA was resuspended in water and quantified by Qubit dsDNA Broad Range kit.

### Slot blot

Slot blot was performed using 1.5 ng of gDNA that was denatured in 400 mM NaOH/10 mM EDTA and blotted onto nitrocellulose membrane (BioRad) in duplicate for dsDNA and 5mC DNA using a slot blot apparatus (BioRad). Membranes were incubated 1 hour at 80°C, blocked with 5% bovine serum albumin (BSA) in TBST (37 mM NaCl, 20 mM Tris pH 7.5, 0.1% Tween 20), and incubated overnight at 4°C in either anti-dsDNA (Abcam, 1:5,000 in 2% BSA in TBST) or anti-5-methyl-cytosine (5mC – Aviva Biosystem clone 33D3, 1:3,000 in 2% BSA in TBST). Membranes were washed in TBST and probed with anti-mouse HRP secondary antibody (Promega; 1:2,000 in 5% BSA in TBST) for 1 hour at room temperature followed by development in ECL (Thermo Fisher Scientific) or Clarity ECL (BioRad). ChemiDoc (BioRad) was used to detect and quantify the chemiluminescent signal. Gel Analyzer (http://www.gelanalyzer.com) was used to perform quantitative densitometric analysis of the signals and ratio between 5mC and dsDNA was plotted for each sample using GraphPad Prism.

### Protein extraction and Western blotting

Frozen tissues were ground using a mortar and pestle cooled with liquid nitrogen and placed in dry ice. 15 mg of tissue powder was used to extract proteins in lysis buffer (20 mM Tris-HCl, 150 mM NaCl, 1% v/v NP-40, 10% v/v glycerol, 2mM EDTA). Protein extraction was performed using a probe sonicator (2 sec pulse, 2 sec pause for 5 min at amplitude 30%) and lysates were cleared by centrifuging at 11,000 × g for 15 min at 4°C and quantified using Qubit reagent (Invitrogen). For preparation of samples for SDS PAGE, 4x Laemmli buffer (BioRad) was added to protein extracts, incubated at 95 °C for 5 minutes and 15 µg of proteins were loaded onto 12.5% denaturating gels, electrophoresed, transferred onto PVDF membranes (BioRad), blocked with 5% w/v powdered milk in TBST buffer (20 mM Tris-HCl, 150 mM NaCl, 0.1% v/v Tween 20, pH 8.0) for 1 hour at room temperature, and incubated overnight at 4°C with anti-H3 (SantaCruz, sc-10809, Rabbit polyclonal, 1:5,000), anti-H3K4me3 (Abcam, ab8580, Rabbit polyclonal, 1:1,000), anti-H3K9me3 (Active Motif, 39161, Rabbit polyclonal, 1:1,000) or anti-H3K27me3 (Active Motif, 61017, Mouse monoclonal, 1:1,000) diluted in blocking buffer. After washing with TBST and incubation for 1 hour with anti-Rabbit IgG HRP Conjugate (Promega, 1:2,000) or anti-Mouse IgG HRP Conjugate (Promega, 1:2,000) diluted in blocking buffer followed by washing in TBST, membranes were visualized using Pierce™ ECL Western Blotting Substrate (Thermo Fisher Scientific) or Clarity ECL substrate (BioRad) on the BioRad ChemiDoc. Immunoblot bands were quantified by densitometry using GelAnalyzer (http://www.gelanalyzer.com).

### RNA-seq

Total RNA extracted as described above was treated by DNAse I for 30 minutes at 37°C followed by RNA purification (RapidOut DNA Removal Kit – Thermo Fisher Scientific). RiboZero was used to remove ribosomal RNA and the remaining sample was used for library preparation according to manufacturer’s instructions (Illumina) from 250 ng of RNA. Libraries were sequenced on NextSeq550 (Illumina) to obtain 150 bp paired-end reads. Raw FASTQ sequenced reads where first assessed for quality using FastQC v0.11.5 (http://www.bioinformatics.babraham.ac.uk/projects/fastqc/). The reads where then passed through Trimmomatic v0.36 [53] for quality trimming and adapter sequence removal, with the parameters (ILLUMINACLIP: trimmomatic_adapter.fa:2:30:10 TRAILING:3 LEADING:3 SLIDINGWINDOW:4:15 MINLEN:36). The dataset from hatchling was also processed with Fastp [54] in order to remove poly-G tails and Novaseq/Nextseq specific artifacts. Following the Fastp quality trimming, the reads were assessed again using FastQC. Quality trimmed reads were used to produce psuedoalignments using Kallisto [55], and the Kallisto quantification was assessed with the --bias flag using the reference *O. bimaculoides* genome (PRJNA270931) and its corresponding annotation. The resulting transcripts per kilobase per million (TPMs) from the pseudo-counts were used for further downstream analysis. Files are available on GEO (accession number: GSE188925).

### Trinotate transcriptome annotation

The quality trimmed reads were aligned to the *O. bimaculoides* genome (PRJNA270931) using HISAT2 [56] with the default parameters and additionally by providing the –dta flag. The resulting SAM alignments were then converted to BAM format and coordinate sorted using SAMtools v1.3.1 [57]. The sorted alignments were processed through Stringtie v1.3.0 [58] for transcriptome quantification to produce a GTF file per sample. The GTFs were then combined using STRINGTIE merge to produce one merged GTF representing the transcriptome for the genome. Finally, Qualimap [59] v2.2.2 was used to generate RNAseq specific QC metrics per sample. Following the transcriptome quantification steps above, the Trinotate [60] pipeline was used to annotate the transcriptome, in addition to the existing reference annotation. The Trinotate steps as detailed in the software’s user manual were followed. Briefly, after transcriptome preparation, BlastP longest ORFs using the longest_orfs protein sequences against the Uniprot database and Pfam domain search using longest ORFs against the Pfam database were integrated into coding region selection using TransDecoder.Predict. In addition, the transcriptome FASTA file was BlastX against the Uniprot database and domain scanned using HMMscan to generate gene to transcript mappings using transIDmapper.pl with the output exported to an SQLite database.

### Reduced Representation Bisulfite Sequencing (RRBS)

RRBS was performed on gDNA extracted from the same 30 dpf hatchling of *O. bimaculoides* used in RNA-seq. Briefly, 1,000 ng of genomic DNA was digested with 200 U of MspI (New England Biolabs) for 24 hours at 37°C. Digested DNA was used for preparing the library as previously described [61], with the exception that adaptors used for multiplexing were purchased separately (Next Multiplex Methylated Adaptors - New England Biolabs). Libraries were size-selected by dual-step purification with Ampure XP Magnetic Beads (Beckman Coulter, Agencourt) to specifically select a range of fragments from 175 to 670 bp, as previously described [62]. Bisulfite conversion was performed with Lightning Methylation Kit (ZYMO Research) by following the manufacturer’s instructions. Libraries were amplified using KAPA HiFi HotStart Uracil+ Taq polymerase (Roche) and purified with Ampure XP Magnetic Beads (Beckman Coulter, Agencourt) before sequencing. Libraries were sequenced using the Illumina Nextseq550 sequencer. Quality control was undertaken using FASTQC (http://www.bioinformatics.babraham.ac.uk/projects/fastqc). Reads were quality trimmed using Trimmomatic [53] to remove low quality reads and adapters. Reads passing quality control were aligned to the reference genome (assembly PRJNA270931, available here: https://groups.oist.jp/molgenu/octopus-genome) using default parameters in Bismark [63], which adopts Bowtie2 as the aligner [64] and calls cytosine methylation at the same time. Fastq files are available on BioSample (accession number: SAMN23139394). RRBS metrics describing lengths of reads, numbers of sequenced and mapped reads and identified citosines were extracted using ‘bismark2report’ function in Bismark (Table S1A), while conversion rates (99.20%) and coverage stats were extracted using ‘processBismarkAln’ and ‘getCoverageStats’ function in methylKit (Table S1B) [65].

### Bioinformatic analysis

RNA-seq data was analyzed as described above and visualized in RStudio (R version 4.0). Heatmaps for transcriptomic profiling were performed by using R package ‘pheatmap’. Clusters were calculated on the hierarchical clustering of dendrogram (Euclidean distance) based on the normalized expression profile (row z-score across different tissues). The number of clusters was determined based on the optimal performance to discriminate the different tissues used for the analysis. For Gene Ontology (GO), GO terms were downloaded from Ensemble Metazoa (BioMart database). GO enrichment analysis was conducted using the GO hypergeometric over-representation test in the ‘ClusterProfiler’ package in R. An adjusted p-value < 0.05 was treated as significant for all analyses. Unique stable transcript IDs were annotated with GO terms from BioMart database and divided by Molecular Function, Biological Process, and Cellular Component terms. Putative members of epigenetic machinery represented in the heatmaps were identified using Trinotate as described above and reported with corresponding human gene symbols. For RRBS analysis on hatchling samples, CpG methylation levels were extracted from Bismarck aligned file with the R package ‘methylKit’ [65]. CpGs covered at least 10 times (and with minimum phred quality score = 20) were included in the analysis. Whole genome bisulfite sequencing (WGBS) data from the Supra and Sub Esophageal brain regions were obtained from public available GEO datasets (GSE141609) [66]. Specifically, Supra (GSM4209498) and Sub (GSM4209499) esophageal brain Bisulfite-seq files (CGmap files) were downloaded and used as source of CpGs methylation analysis. To compare WGBS data with RRBS, CGmap files were processed as follow: CGmap files were filtered for methylation on CpGs context only (based on column 4 representing the context (CG/CHG/CHH)); strand direction information was obtained converting C into + and G into – (based on column 2 representing the nucleotide on Watson (+) strand); percentage of methylation was obtained multiplying by 100 the methylation-level (from 0 to 1; in 0 to 100 values) (based on column 6 methylation-level). CpGs with methylation levels below 20% were treated as unmethylated and above 80% were considered as methylated. Genomic element annotation and metaplots of CpGs were performed with R package ‘genomation’. Repetitive Elements (RE) were identified using the Repeat Masker annotation on the reference genome (assembly PRJNA270931, available here: https://groups.oist.jp/molgenu/octopus-genome), and manually curated to group the classes of transposons (DNA, LTR, LINE, SINE) and non-transposons (Satellite, Simple Repeats, Other RE). Since sequences represented (transcripts or RE) are of unequal length, metaplots were divided into 30 or 15 equal bins (based on the average length of the sequences analyzed). Lines in metaplots represent the winsorized mean values (1–99 percentile) for each bin, and blue shades represent dispersion of 95 percent confidence interval for the mean. Heatmaps of DNA methylation pattern were performed with R package ‘EnrichedHeatmap” on the full-length transcripts and 2,000 bp upstream and downstream. DNA methylation average of gene bodies for each transcript was divided into 4 clusters based on kmeans from R package ‘stats’. Number of clusters was optimized at 4 based on the ability to discriminate between the distinct patterns of DNA methylation. Center of kmeans from Pattern 1 to 4 are respectively 71.06, 49.11, 28.59, 1.31. Subsequent analyses on methylation pattern were performed using R package ‘ggplot’ for violin plots, ‘pheatmap’ for expression profiling, and ‘ClusterProfiler’ for GO enrichment analysis. All code used for this study is available on Github (https://github.com/SadlerEdepli-NYUAD).

### Phylogenomic Pipeline

Human sequences for genes encoding DNA methylation and histone modification factors were collected from Uniprot (October 5, 2021) to generate reference gene sets (RGS) per family (Table S2). Project databases, Metazoa50 and Metazoa19 (a subsample of Metazoa50), representing 50 and 19 animal and unicellular outgroup genomes, respectively, were generated per genome. Genomes were sourced from Ensembl, NCBI RefSeq, or other public databases (Table S3), and species and data set selection was based on representation of major clades and phyla and BUSCO-based evaluations of genome quality (BUSCO; Metazoa10) [67]. Genome gene models were filtered to retain single longest protein isoform per coding gene locus. Fasta file headers were standardized per genome based on NCBI Taxonomy (January 2021) details per species and on common names as provided in Wikipedia (October 2021) (Table S2). Blastp databases were generated per genome using NCBI Blast+ (version 2.6.0) Makeblastdb. Metazoa50 genomes and RGS sequences were domain annotated using Interproscan Pfam (version 5.48-83.0) [68].

Human reference gene sets were blasted against Metazoa19 and Metazoa50 Blast databases per gene family or superfamily using NCBI Blast+ Blastp (e-5 cutoff). All hits were then blasted against the human genome and top hits to any human RGS sequence retained to form the candidate gene set (CGS), which were combined with RGS sequences to form a final gene set. Sequences were aligned in MAFFT (linsi command; version 7.487)[69], alignments were trimmed in ClipKIT (super gappy; version 1.1.3) [70], and maximum likelihood trees were built in IQTREE2 (with ModelFinder; version 2.1.2) [71]. Alignments and trees were visually assessed in Geneious (version 2021.1.1) and in FigTree (version 1.4.4 and iTOL (version 6) [72], respectively. Sequences residing on UHRF1, DNMT1, SETD1B, SETDB1, EZH2, KAT2A, and HDAC8 branches within a larger superfamily tree were collected and then aligned, trimmed, and a tree built for just the family to improve branching structure and support relative to the superfamily tree. Octopus sequences came from the NCBI RefSeq genome and were mapped to the Ensembl genome based on top Blastp hit to match sequence identifier to those used in transcriptome analyses. Trees were color annotated in FigTree and rooted on sponge or a unicellular outgroup or else left unrooted.

### Protein alignment

Protein sequences were downloaded from the UniProt database (https://www.uniprot.org) as FASTA formatted files and alignments were performed using ClustalOmega with multiple sequence alignment program [73]. Output alignment files generated in a ClustalW format with character counts were reformatted and colored based on amino acid residue identity using MView (https://www.ebi.ac.uk/Tools/msa/mview/). Protein sequences of interest were processed with phmmer [74] and queried against HMM target databases using profile hidden Markov models. Sequences matches were calculated by multiple factors, grouped by Pfam domains and homology sequence probability were represented by Bit score.

### Swiss-Expasy 3D Model

To build protein homology models we used the Automated Mode (https://swissmodel.expasy.org/docs/help) of the SWISS-MODEL server homology modeling pipeline [75]. In brief, homology modeling proceeds through four main steps: (i) alignment of target sequence and identification of structural templates by BLAST and HHblits; (ii) alignment and sorting of target–template structures based on Global Model Quality Estimation (GMQE) and Quaternary Structure Quality Estimation (QSQE); (iii) model-building relying on ProMod3 [76] and OpenStructure comparative modelling engine [77]; and (iv) model quality evaluation using GMQE estimation score, and another composite estimator QMEAN [78]. SWISS-MODEL Structure comparison tool was used to perform super-positioning of newly computed 3D models of *O. bimaculoides* proteins and published structures of mouse DNMT1 (PDB ID 4da4.1) and UHRF1 (PDB ID 2zke.1).

### Statistical analysis

All experiments were carried out on at least 3 biological replicates, except where noted. For slot blot analysis, technical replicates were also included. The number of replicates for each experiment is indicated in each figure. Methods to evaluate statistical significance include unpaired parametric one-way ANOVA test adjusted with Tukey’s multiple comparisons test or unpaired non-parametric Kruskal-Wallis test adjusted with Dunn’s multiple comparisons test. Tests used are indicated in each graph. All plots were generated in GraphPad Prism 9 or RStudio (R version 4.0). Statistical analysis was performed in GraphPad Prism 9.

## Results

### Tissue specific gene clusters identified in octopus

We extended previously published RNA-seq datasets obtained from 11 different adult tissues (Figure 1A; [22]) by generating new RNA-seq datasets from the first left (L1) arm of an adult male octopus at distal, medial and proximal locations (between 1.5 and 5.5 cm from the arm tip, Figure S1A-B), and one 30 day old hatchling (Figure S1B-C). We combined these datasets and found that of 40,327 transcripts annotated in the *O. bimaculoides* genome, 38,232 were expressed in at least one sample (TPM > 0). Trinotate [79] was used to better annotate the *O. bimaculoides* genome and assigned putative gene names and putative UniProt entry names to 23,879 transcripts (59%) identified in the datasets analyzed (Table S4).

**Figure 1.**
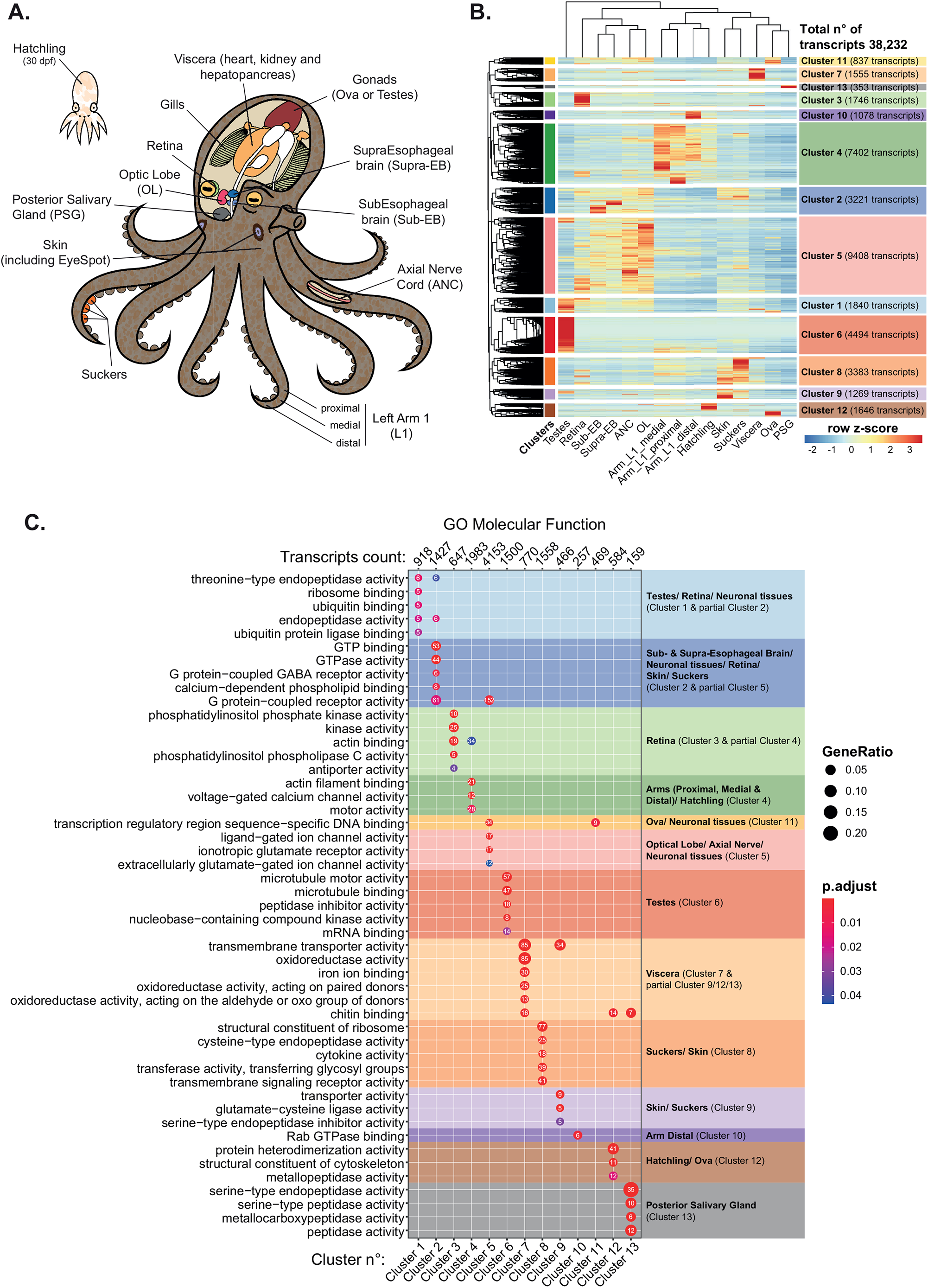
Expression profiling identifies tissue specific gene-clusters with distinct functional annotation. **A**. Schematic representation of *O. bimaculoides* anatomy highlighting the tissues analyzed in this study. **B**. Heatmap of the expression profile of 13 different tissues (including 12 different tissues derived from adult animals and one whole-body hatchling). Rows are divided into 13 clusters calculated on the hierarchical clustering of dendrogram (Euclidean distance). **C**. Gene Ontology (GO) of genes in each cluster. Dot size represents gene ratio between observed and expected transcripts in each GO category (with padj < 0.05), dot colors represent adjusted p-value and numbers inside dots indicate number of transcripts for each GO term in each cluster.

Optimized hierarchical clustering of the 38,232 expressed transcripts generated 13 clusters that define specific transcriptomic profiles across these tissues (Figure 1B and Table S5). Some clusters were dominated by transcripts that were nearly exclusively expressed in a single tissue, such as Cluster 6 (testes), whereas other clusters were defined by transcripts expressed in multiple tissues that are functionally related, such as Cluster 5 (tissues with neuronal functions). Interestingly, the distal arm segment was characterized by a distinct set of transcripts in Cluster 10 that were not highly enriched in any other tissue (Figure 1B), potentially reflecting the sensory and regenerative capacity of this structure [36, 41, 80].

Gene Ontology (GO) analysis for Molecular Function (Figure 1C), Biological Process (Figure S2A) and Cellular Component (Figure S2B) revealed that each cluster was enriched for transcripts encoding proteins with shared functions. For example, microtubule binding, motor activity, and cilium movement terms which are prominent for sperm activity characterized genes expressed in testes (i.e. Cluster 6; Figure 1C and S2A). G-protein coupled receptor activity, GTP binding, glutamate receptor activity, and signaling pathways and others critical for neuronal function and were enriched in Clusters 2 and 5, (brain and nervous tissues; Figure 1C and S2A). Cluster 4, which was enriched in transcripts expressed in the arm and hatchling, included terms involved in motor activity, actin filament binding, calcium-ion transmembrane transport, and myosin complex (Figure 1C and S2A-B), reflecting contractile activities of muscles. These findings make predictions about genes regulating cell identity and key processes in *O. bimaculoides* tissues and suggest that complex regulatory mechanisms, potentially based in the epigenome, regulate these gene expression profiles.

### DNA methylation machinery is conserved and differentially expressed in octopus

We characterized DNMT and UHRF gene family evolution in animals using a phylogenomic pipeline we designed. Human DNMT1, TRDMT1 (DNMT2), DNMT3A, DNMT3B, and DNMT3L and human UHRF1 and UHRF2 were compared against Metazoa19 and Metazoa50 genomes. We identified DNMT1 (Ensembl Transcript ID: Ocbimv22034501m; NCBI gene symbol: LOC106877272; Figure 2A and S3A) and UHRF1 (Ensembl Transcript ID: Ocbimv22021185m; NCBI gene symbol: LOC106874972; Figure 2B and S3B) homologs in *O. bimaculoides* and across a range of animal species. Both DNMT1 and UHRF1 are absent in the three unicellular outgroups examined (choanoflagellate, *Mongosiga brevicollis*, the flilasterean, *Capsaspora owczarzaki* and the ichthyosporean, *Sphaeroforma artica*), but highly conserved in most animals and were likely present in the last common ancestor, with a number of independent losses in diverse lineages. DNMT1 was lost in 11 out of 47 Metazoa50 animal species, including the fruit fly *Drosophila melanogaster* (but was present in two other arthropods), and nematodes. UHRF losses were detected for 12 out of 47 Metazoa50 animal species, which match losses of DNMT1 except for the sea squirt, *Ciona savignyi*, which is predicted to have low level of DNA methylation [81]. Several species or clades also exhibited expansions to two or three copies for both genes (Figure 2A-B and S3A-B).

**Figure 2.**
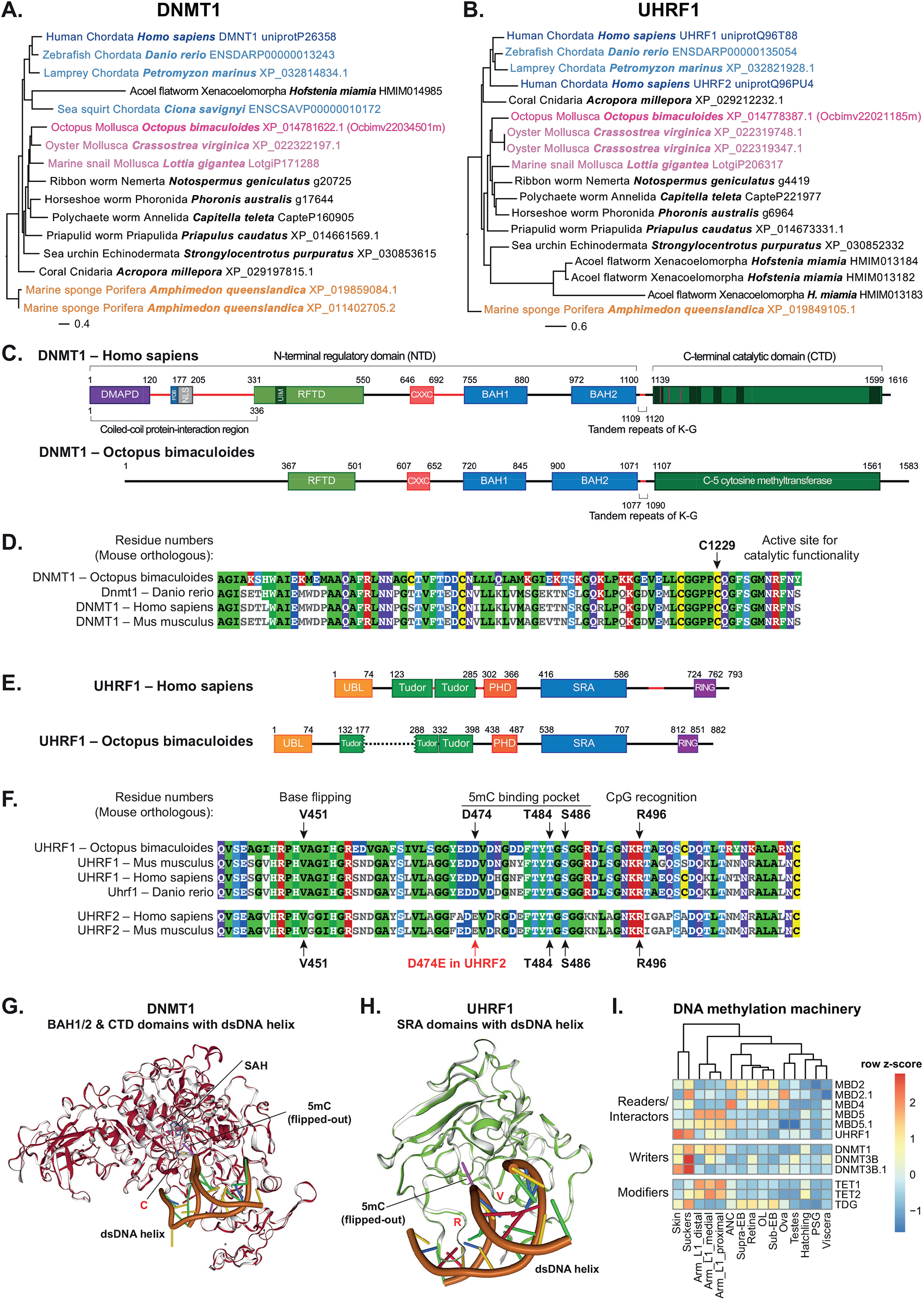
DNA methylation machinery is conserved and differentially expressed across different tissues. Phylogenetic trees of **A**. DNMT1 and **B**. UHRF1 in a representative subset of 19 metazoan and outgroup species. **C**. DNMT1 domain structure in *H. sapiens* and *O. bimaculoides*. Numbers indicate amino acid residues for each species. **D**. Alignment of the C-terminal Catalytic Domain (CTD) of *O. bimaculoides* to *M. musculus, H. sapiens* and *D. rerio* shows that the major residue needed for DNMT1 catalytic function is highly conserved among the species. Residue functionality was assigned based on the mouse DMNT1 ortholog. **E**. Domain structure of UHRF1 in *H. sapiens* and *O. bimaculoides*. **F**. Alignment of the *O. bimaculoides* SRA domain to *M. musculus, H. sapiens* and *D. rerio* shows that all major residues needed for UHRF1 functionality are conserved among the species. Residue functionality is based on the mouse UHRF1 ortholog [24, 26]. Alignment to UHRF2 shows no conservation of the critical residues between the SRA domain in *O. bimaculoides* and UHRF2 in *H. sapiens* and *M. musculus*. **G**. Structural superposition of the 3D structure of BAH1, BAH2, and CTD domains in *M. musculus* (grey) with the 3D model of the same domains in *O. bimaculoides* (red). DNA is represented in brown and critical residues for 5mC deposition are highlighted in red. **H**. Structural superposition of the 3D structure of the SRA domain in *M. musculus* (grey) with the 3D model of the same domain in *O. bimaculoides* (green). DNA is represented in brown and residues critical for CpGs recognition and 5mC base flipping are highlighted in red. **I**. Expression profiles of DNA methylation machinery. Gene names are extracted from Trinotate (Table S6).

The DNMT1 transcript (hereafter termed DNMT1_OCTBM) encodes a protein that retains the domain structure (Figure 2C) and key amino acid residues (Figure 2D) that are necessary for its methyltransferase function. HMMER analysis of DNMT1_OCTBM to identify sequence homology in the animal reference protein database showed that the C-5 cytosine methyl-transferase domain (Pfam ID: DNA_methylase), which carries out the catalytic function of DNMT1, had the highest homology score and overall, DNMT1_OCTBM had over 50% of identity with vertebrate DNMT1s (Figure S4A-B). We conclude that DNMT1_OCTBM has the necessary features to function as a DNA methyltransferase and to interact with UHRF1.

UHRF1 and UHRF2 family members in mammals, but there is only 1 family member present in mollusks (Figure S3B and [18]). The protein encoded by Ocbimv22021185m (hereafter termed UHRF1_OCTBM) retains features of the mammalian protein, including tandem Tudor and PHD domains that read the histone code and the SRA domain, which binds hemimethylated DNA and facilitates DNMT1 access to CpGs (Figure 2E) [23-26, 30, 31, 82, 83]. The SRA domain of UHRF1_OCTBM has highest homology compared to vertebrate homologs (Figure S4C-D), with key residues involved in hemi-methylated DNA recognition and base flipping [24, 26] completely conserved (Figure 2F). There was much lower homology between UHRF1_OCTBM and UHRF2, and the 5mC binding pocket (residue D474E), the base flipping motif (HVAG thumb loop) and the CpG recognition site (NKRT finger loop) [25] were not conserved (Figure 2F). Another octopus transcript (Ocbimv22020196m; termed YDG-OCTBM), encoded a shorter protein containing a domain similar to the SRA but had low homology to vertebrate UHRF1 or UHRF2 (Figure S4E-F) and lacked key residues for CpG recognition found in both UHRF1 and UHRF2 (Figure S4G). This is further supported by Swiss-Expasy 3D modeling [75] which reveal highly conserved 3D structures for DNMT1_OCTBM, with a near identical structures to the mouse protein (Figure 2G, S5A-B), including DNA double helix correctly located, the 5mC exposed near the C1194 (C1229 in mouse [84, 85]) and S-Adenosyl-L-methionine properly positioned (Figure 3G, S5A-B). Furthermore, the SRA domain of UHRF1_OCTBM aligned to the mouse domain, with the loops necessary to bind DNA, to recognize CpGs (R612 and R496 in mouse [24, 26]) and to mediate base-flipping (V567 and V451 in mouse) all properly positioned (Figure 2H, S5C-D). This indicates that Ocbimv22034501m encodes the closest DNMT1 homolog and Ocbimv22021185m encodes the UHRF1 ortholog in *O. bimaculoides* and that the encoded proteins retain functional capacity to interact as a complex, for UHRF1 to recognize hemi-methylated DNA and for DNMT1 to methylated CpGs.

**Figure 3.**
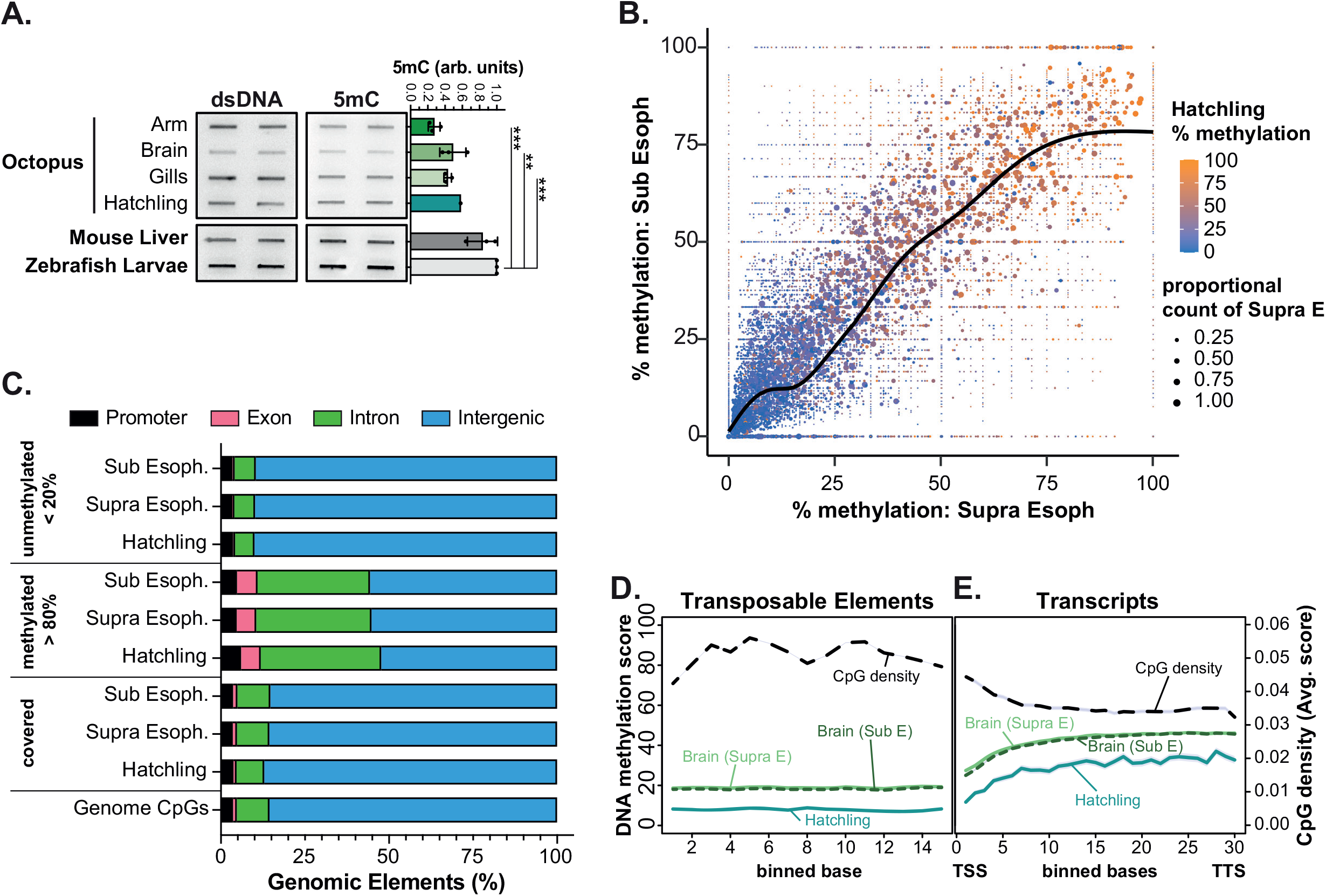
DNA methylation is enriched in a subset of gene bodies in *O. bimaculoides*. **A**. Slot blot of gDNA extracted from the distal arm tip (1.5 cm of right arm 2), brain (optical lobe), and gills of one representative *O. bimaculoides* adult animal, and whole 30 day od hatchling (same biological sample used for RNA-seq and RRBS). gDNA extracted from one representative male mouse adult liver and one representative pool of whole 5 dpf zebrafish larvae was blotted on the same membrane for comparison. The 5mC signal was normalized to double stranded DNA (dsDNA) and then each sample was normalized to levels zebrafish larvae. Each dot represents one biological replicate. p-value were calculated by unpaired parametric one-way ANOVA test adjusted with Tukey’s multiple comparisons test. Adjusted p-value are indicated as *** < 0.001, ** < 0.01. Comparisons other than with zebrafish or mouse resulted in not significant adjusted p-values > 0.05 (not indicated in the graph). **B**. Scatter plot of DNA methylation levels of common CpGs across Supra Esophageal (Supra E) and Sub Esophageal (Sub E) brain. Dot color represents DNA methylation levels in hatchling and size of the dots indicates scaled proportion of CpGs represented by each dot. **C**. Genomic annotation of CpGs contained in *O. bimaculoides* genome, and of those covered by WGBS in brain and RRBS in hatchling. CpGs were divided into methylated (> 80%) and not methylated (< 20%) CpGs and relative genomic annotation was performed. **D**. Metaplot represents DNA methylation levels of transposable elements in Supra E and Sub E brain and hatchling. CpG density of octopus genome is represented for the same region. Each region is divided in 15 bins. **E**. Metaplot represents the DNA methylation levels in Supra E and Sub E brain and hatchling for full-length transcripts. CpG density of octopus genome has been represented for the same region. Each transcript has been divided in 30 bins from Transcription Star Site (TSS) to Transcription Termination Site (TTS).

UHRF1 and DNMT1 function as a complex, and are typically co-expressed in cells that are actively proliferating. We found this to be the case in octopus tissues, with highest levels of both genes expressed in skin, suckers and arms (Figure 2I). This is consistent with the idea that these cells experience high turnover are expected to have proliferating cells [80]. Interestingly, we found that most genes encoding factors that read (methyl binding proteins; MBD), write (DNMTs) and erase (ten-eleven translocation; TET) DNA methylation in octopus were expressed at low levels in most tissues except for arms, suckers and skin (Figure 2I, Table S6). The relevance of this pattern is not yet known.

### DNA methylation is enriched in the bodies of a subset of genes in *O. bimaculoides*

Reports of the presence of DNA methylation in octopus [16, 51, 52] and other mollusks [13, 15, 20] indicate that the DNA methylation machinery is functional in these animals. To test this directly, we used slot blot analysis to detect bulk 5mC levels on genomic DNA (gDNA) extracted from arm tip, brain (optic lobe) and gills of 3 different *O. bimaculoides* adults and a hatchling. We found relatively equivalent levels in all tissues, albeit at less than half of what was detected in mouse liver or zebrafish larvae (Figure 3A).

We next examined the genome-wide distribution of methylated and unmethylated CpGs by performing RRBS on gDNA isolated from a 30 dpf hatchling and by analyzing previously published WGBS datasets from 2 regions of the adult brain (Supra Esophageal - Supra E; and Sub Esophageal - Sub E) [16]. There are 80,660,576 CpGs in the *O. bimaculoides* genome, considering both DNA strands. RRBS profiling of the hatchling genome covered 2.98% of these (2,403,266 CpGs) and WGBS of the Supra E and Sub E covered 80.39% (64,845,686 CpGs) and 58.33% (47,053,504 CpGs), respectively (Figure S6A). In all samples, the vast majority of CpGs had methylation levels categorized as not methylated (<20% methylation) and CpGs categorized as methylated (>80% methylated) was low, ranging from 3.63% in hatchlings and 7.53% in the Sub E brain (Figure S6A-B). Approximately 7% of CpGs showed an intermediate pattern of methylation (between 20- 80%) which could reflect tissue heterogeneity or a stochastic pattern of methylation on these CpGs. The methylation pattern (Figure S6B) is in marked contrast to the bimodal pattern found in vertebrates and some species of sponge [17, 18], where majority of CpGs are methylated and the remaining are unmethylated. These findings indicate that the octopus methylome resembles other invertebrates [3, 8, 12] including another mollusk, the Japanese oyster *Crassostrea gigas* [3, 14], but is distinct from invertebrate model organisms which lack methylation entirely.

DNA methylation serves to repress transposon expression in vertebrates, and therefore the bulk of methylated CpGs are found in intergenic regions, with very little tissue-specific variation [3, 86, 87]. To investigate if there was a tissue specific CpG methylation pattern in octopus, we identified 1,425,557 CpGs that were commonly detected in the 2 brain and hatchling methylome datasets and then plotted the methylation levels on each CpG in the two brain samples and overlaid the methylation levels from the hatchling (Figure 3B). This showed a linear correlation between brain samples and that nearly all CpGs that were either unmethylated or highly methylated in brain had the same pattern in hatchling (Figure 3B), indicating little tissue-specific variation.

The scaffold nature of the *O. bimaculoides* genome [22] makes it difficult to accurately approximate the proportion that is intergenic compared to genic, but clearly the vast majority is intergenic, which is also where over 85% of CpGs reside (Figure 3C). These intergenic CpGs are comparatively depleted of methylation and there is a strong enrichment of methylated CpGs (>80%) detected in introns and exons (Figure 3C). Moreover, while there was high CpG density across the length of TEs, methylation levels were consistently low (Figure 3D), regardless of the TE class (Figure S6C). In contrast, DNA methylation was comparatively higher in gene bodies, with the average methylation level across gene bodies did not exceed 50% in any tissue (Figure 3E). Although we cannot exclude that our finding of low methylation levels across TEs could be attributed to their poor annotation, our data indicate that DNA methylation is not a major epigenetic mechanism of TE repression in octopus. We speculate that in cephalopods, like in other species with low DNA methylation levels, gene bodies are the main target of methylation [12].

The 5mC is a binary mark, so that each residue in individual cells can be either methylated or unmethylated. Therefore, we reasoned that the intermediate methylation pattern observed across all gene bodies represented heterogeneity in the CpG methylation pattern of genes. We tested this in the Supra E brain sample since it was the sample with the highest amount of covered CpGs (Figure S6A). Using k-means clustering of genes based on CpG methylation levels averaged for each transcript identified 4 distinct Methylation Patterns (Figure 4A-B): Pattern 1 had the fewest number of genes (5,482) and was characterized by having high CpG methylation across the entire gene body and promoter; this was distinguished from genes in Pattern 2 (8,972), which had high methylation across the gene body and downstream of the transcription termination site (TTS) but lacked methylation at the transcript start site (TSS) and promoter. Genes in Pattern 3 (6,545) were characterized by having unmethylated CpGs in the first third of the gene body but higher methylation in the 3’ end of the gene and around the TTS. Pattern 4 was the inverse of Pattern 1, as it had the largest number of genes (17,586) and virtually no methylated CpGs (Figure 4A and Table S7). Strikingly, these Methylation Patterns were the same in the Sub E and hatchling samples (Figure 4B, Figure S7A-B), indicating that the pattern of DNA methylation on gene is set by a cellular or genomic feature that is consistent across tissues.

**Figure 4.**
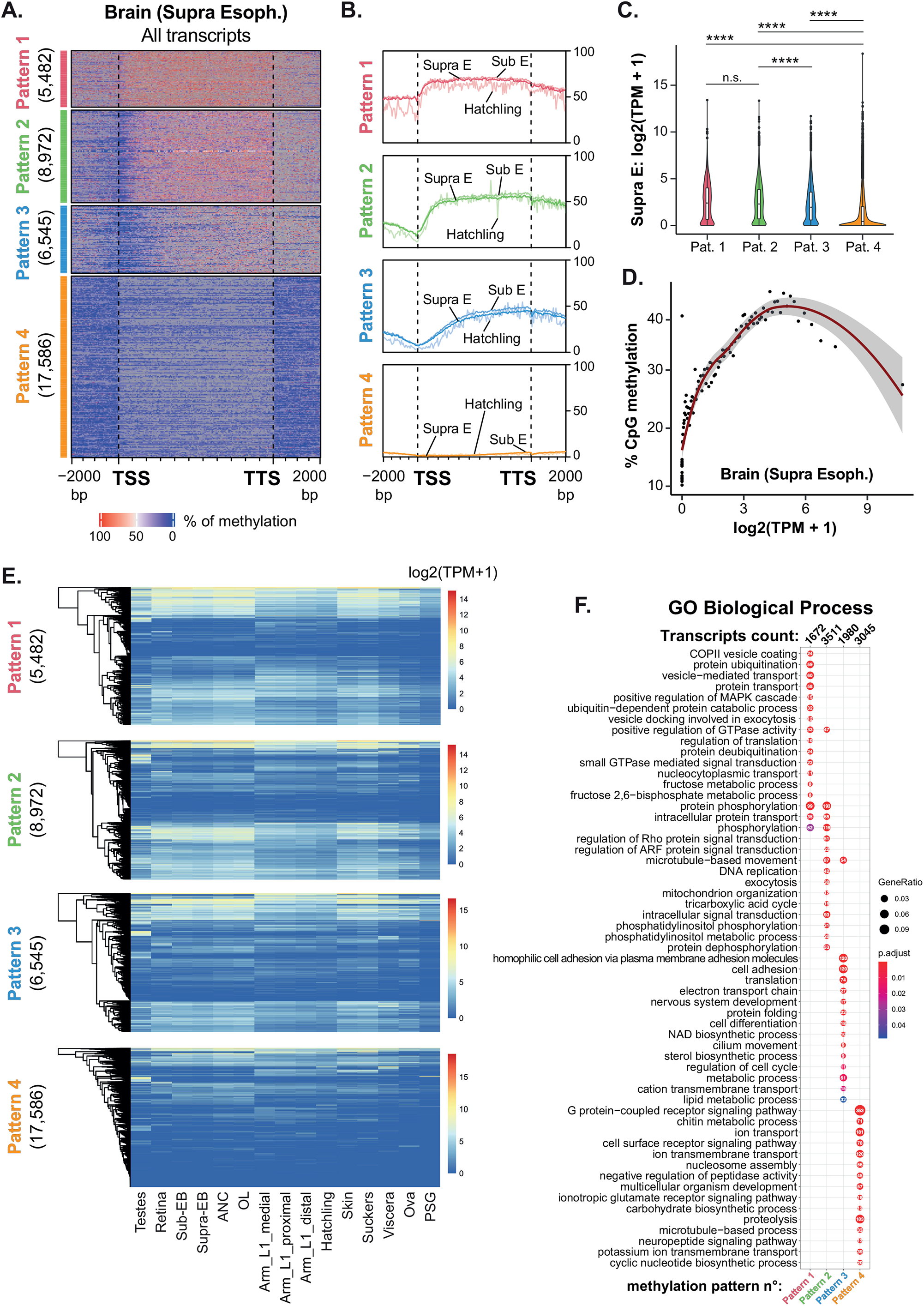
DNA methylation on gene bodies correlates with gene expression. **A**. Heatmap of DNA methylation levels detected in Supra E brain samples across full-length transcripts and 2000 bp upstream and downstream of start site. All transcripts have been divided in 4 clusters by k-means based on the DNA methylation average across each transcript. **B**. Line plot represents average of DNA methylation of Supra E and Sub E brain and hatchling for each cluster defined in the heatmap on Supra E sample. **C**. Violin plot displays the distribution of transcript expression values (as log2(TPM+1)) for each methylation pattern in Supra E brain samples. p-values were calculated by unpaired non-parametric Kruskal-Wallis test adjusted with Dunn’s multiple comparisons test. **** indicates adjusted p-value < 0.0001, and n.s. indicates not significant adjusted p-values > 0.9999. **D**. Overall DNA methylation of transcripts (TSS to TTS of each transcript) in Supra E brain is regressed against transcripts expression (log2(TPM+1)). Transcripts were grouped by percentile of expression values and each dot represents the average value of DNA methylation for each percentile. **E**. The expression profile (as log2(TPM+1)) of transcripts categorized in each methylation pattern have been represented across the 13 different tissues. Column clustering is supervised according to the sample order established in Fig 1B. Row clustering is calculated on the hierarchical clustering of dendrogram (Euclidean distance). **F**. Gene Ontology (GO Biological Process) annotation of transcripts in each methylation pattern. Dot size represents gene ratio between observed and expected transcripts in each GO category (with padj < 0.05), dot colors represent adjusted p-value and numbers inside the dots indicate number of transcripts for each GO term.

We determine the relationship between gene body methylation and expression by plotting the expression (log2(TPM+1)) of all the transcripts in each Methylation Pattern. The highly methylated genes (Patterns 1-2) were expressed at the highest levels, while unmethylated genes (Pattern 4) were expressed at the lowest levels in the Supra E samples (Figure 4C). To examine the direct correlation between methylation level and expression, transcripts in the Supra E sample were grouped into percentiles based on expression levels (log2(TPM+1)) and then calculated the mean methylation level for all genes in each percentile (Figure 4D). This revealed a direct correlation between DNA methylation and expression for genes at the mid-range of expression levels (1.5 to 4.5 log2(TPM+1)) but there was low or no correlation between methylation levels and expression in genes that had low to no DNA methylation (Figure 4D). This correlation was also found in Sub E and hatchling samples (Figure S7C). Strikingly, a subset of genes in Patterns 1 and 2 were expressed at high levels in all tissues, whereas genes in Pattern 4 were either silenced or expressed at more variable levels across tissues. This suggests that hypermethylation of gene bodies serves to promote expression of genes that are sustained at high levels in all tissues and that hypomethylated genes are silenced.

GO analysis showed that specific gene functions were enriched in each Methylation Pattern. Trafficking, signaling, translation and metabolism characterized genes in Patterns 1-2 and the unmethylated genes in Pattern 4 participated in metabolism, processes that occur at the cell surface, or are related to neuronal functions (Figure 4F). This indicates that the methylation pattern on gene bodies, or lack thereof, defined genes that play roles in similar biological processes. It also suggests that the genes in Pattern 1 may be involved in processes that are required by all cells, such as protein secretion, metabolism and translation.

Studies in other invertebrates suggested a correlation between gene body methylation, gene length, and age [14]. Transcripts in each *O. bimaculoides* Methylation Pattern had a large range of average lengths (from TSS to TTS), with the highest variability in fully methylated transcripts (Pattern 1), with slightly shorter genes in Pattern 4 and slightly longer, on average transcripts in Pattern 2 and 3 (Figure S7D). In summary, this analysis shows that there is a similar pattern of DNA methylation on genes across tissues that positively correlates with gene expression, and genes with distinct functions are marked by similar methylation patterns.

### Conserved epigenetic machinery elicits a dynamic pattern of histone modifications in octopus tissues

H3K9me3 and H3K27me3 are highly conserved marks of repressed chromatin, while H3K4me3 marks actively transcribed genes and histone acetylation serves to open chromatin. We investigated the conservation of some well-studied enzymes that write these marks. There was a many-to-one human-octopus ortholog of the H3K4me3 histone methyl transferase (SETD1B in humans; Figure 5A) and a many-to-one human-octopus ortholog of the H3K9me3 histone methyl transferase (SETDB1 in humans; Figure 5B); both showing high conservation across metazoans. A many-to-one human-octopus ortholog of the H3K27me3 methyltransferases (human EZH2) is present in octopus and conserved across species (Figure S8A). Representative examples from the histone acetylation modifying enzymes KAT2A and HDAC8 revealed a many-to-one human-octopus ortholog of KAT2A and a one-to-one human-octopus ortholog of HDAC8 in octopus and conserved in other animals (Figure S8B-C).

**Figure 5.**
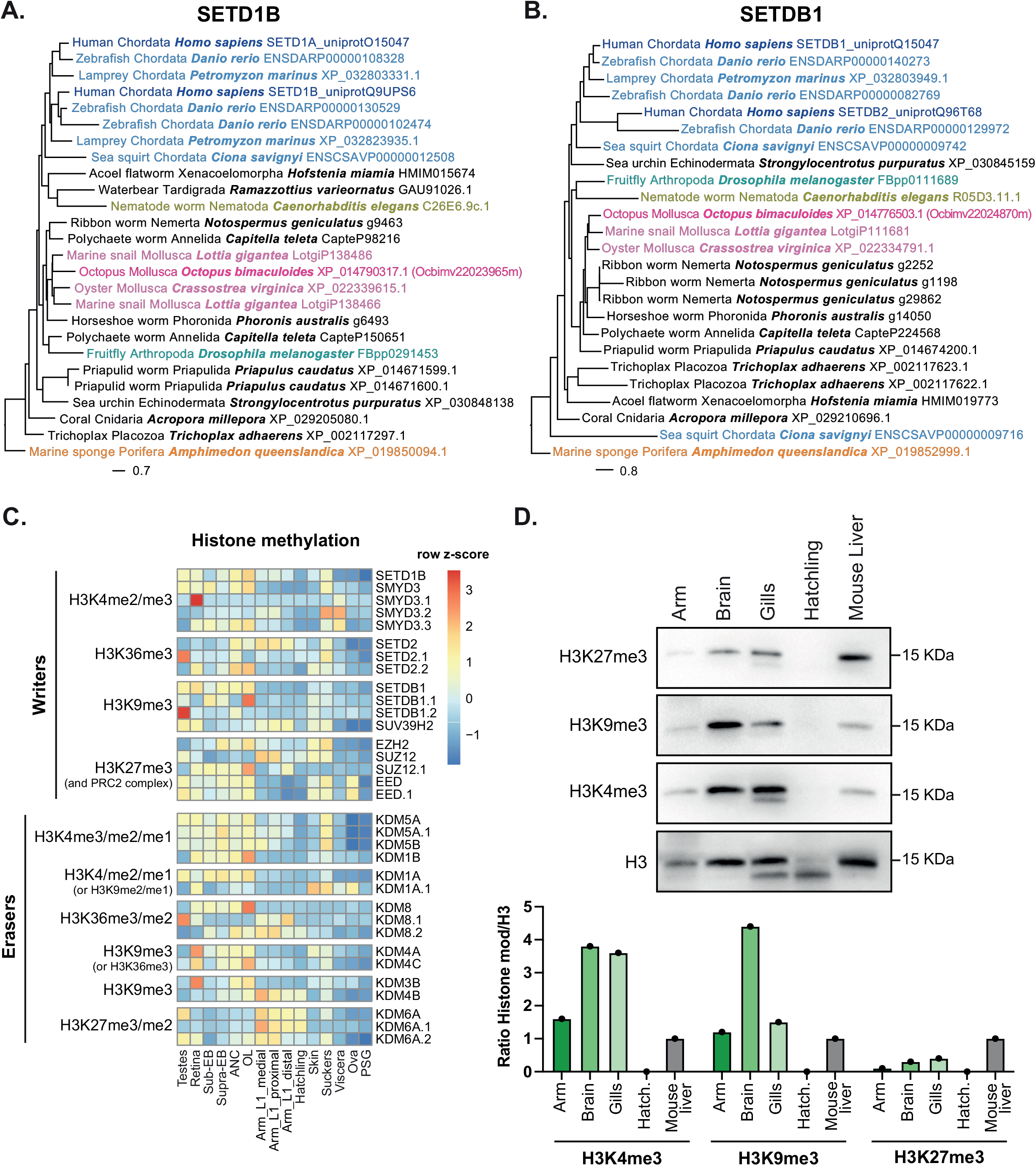
Conserved epigenetic machinery elicits a dynamic pattern of histone modifications in different octopus tissues. **A**. Phylogenetic tree of SETD1B, responsible for H3K4me3 deposition, in a representative subset of 19 metazoan and outgroup species. **B**. Phylogenetic tree of SETDB1, responsible for H3K9me3 deposition, in a representative subset of 19 metazoan and outgroup species. **C**. Heatmap of the main histone methylation factors, methyltransferase (writers) and de-methylases (erasers). **D**. Western blot of histone H3 and relative modification (H3K4me3, H3K9me3, H3K27me3) performed on arm (1.5 cm of Right Arm 2), brain (optical lobe), gills and hatchling of *O. bimaculoides*, in comparison with quiescent mouse adult liver. Quantification of each sample measured by western blot was normalized to histone H3 and each histone mark was normalized to the relative mark in mouse liver as control. Each dot represents one biological replicate.

We next examined the expression profile of these genes and others categorized by Trinotate identification (Table S6) as regulators of histone methylation (Figure 5C and 9A) and acetylation (Figure 9B). A tissue-specific expression profile was identified for most of these genes, with the testes and neuronal tissue expressing the highest levels of the H3K4 and H3K9 methyltransferases as well as high levels of the H3K4 demethylases (Figure 5C, Table S6), and genes that erase H3K27me2/3 were higher in the arms and hatchlings. Interestingly, expression analysis of all histone methyltransferase clearly show that specific subsets of HMTs characterize a particular tissue (Figure S9A). For instance, several of the enzymes that remove methylation from H3K4 (KDM5A/B) are enriched in testes, retina, brain and suckers, while skin and suckers are enriched for the demethylases that act on H3K9 or H3K36 (i.e. KDM4A/C; Figure 5C).

We determine if these enzymes were functionally active in *O. bimaculoides* using Western blotting for H3K27me3, H3K9me3 and H3K4me3 in the arm, brain, gills, and hatchling. This showed a distinct patterns in each tissue: H3K4me3 was detected at high levels in brain and gills and lower levels in the arm, while H3K9me3 was higher in brain compared to gills and arm (Figure 5D). H3K27me3 showed comparable level in brain and gills and lower in arm, albeit overall lower level in all samples compared to H3K4me3 and H3K9me3. The low levels of all histone modifications investigated in hatchling could reflect the low level of expression of all histone methyltransferases in this sample. Although the complex signature of gene expression makes it difficult to predict which histone modification marks prevail in each tissue, these data suggest histone code plays a prominent role in regulating tissue identity.

### DNA methylation and histone marks are present in squid and bobtail squid

To determine if the epigenetic modifications identified in *O. bimaculoides* were conserved in other cephalopods, we analyzed tissues from *D. pealeii* (longfin inshore squid) and *E. berryi* (bobtail squid) for 5mC levels by slot blot H3K27me3, H3K9me3 and H3K4me3 by Western blot. 5mC levels in *D. pealeii* and *E. berryi* tissues were similar to those found in *O. bimaculoides* and markedly lower than levels in mouse or zebrafish (Figure 6A-B). Histone marks were detected in arm tissue from both squid and bobtail squid (Figure 6C). Interestingly, H3K27me3 and H3K4me3 showed comparable levels to the mouse liver, while H3K9me3 was high in squid and bobtail squid arms compared to mouse tissue (Figure 6C-D). Therefore, the mechanisms that regulate the 5mC and histone methylation are conserved in cephalopods.

**Figure 6.**
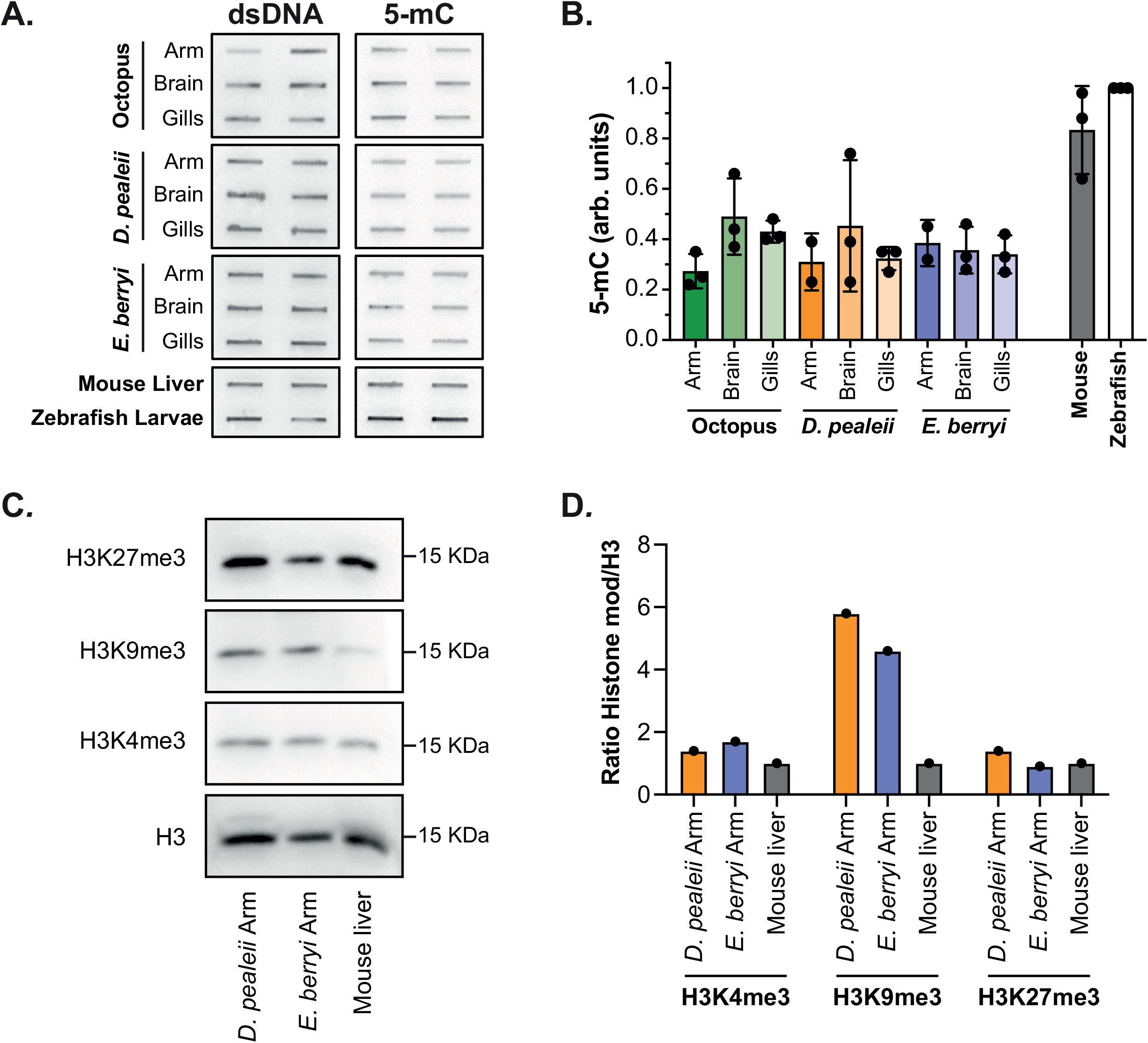
DNA methylation and histone marks are present in squid, and bobtail squid. **A**. Slot blot detecting 5mC on gDNA extracted from arm (2-3 cm of Right Arm 2), brain (optical lobe), gills of one representative animal of *D. pealeii* and *E. berryi*, the analogous tissues of *O. bimaculoides* adults and a 30 dpf hatchling, one representative male mouse adult liver and one representative pool of 5 dpf larvae of zebrafish. **B**. Quantification of 5-mC measured by slot blot was normalized to double stranded DNA (dsDNA) and each sample was normalized to zebrafish larvae as control. Each dot represents one biological replicate. **C**. Western blot of histone H3 and relative modification (H3K4me3, H3K9me3, H3K27me3) performed on an arm of *D. pealeii* and *E. berryi*, in comparison with quiescent mouse adult liver. **D**. Quantification of each sample measured by Western blot was normalized to histone H3 and each histone mark was normalized to the relative mark in mouse liver as control. Each dot represents one biological replicate.

## Discussion

The role of epigenetics in cell identity, plasticity, development, and disease is one of the most investigated aspects of biology that is heavily bias towards vertebrate models. This study identifies new features of the epigenome in cephalopods including DNA and histone methylation. We show that DNMT1 and UHRF1 are evolutionary conserved and identify critical residues necessary for their distinct functions identical in human and octopus homologs. Interestingly, in most species, DNMT1 and UHRF1 evolved together with DNA methylation, so that the presence of these genes predicts the presence of CpG methylation [3, 16-18].

We demonstrate functionality of these factors and those that methylate histone tails across multiple tissues in multiple cephalopod tissues and species. Less than 10% of all CpGs are methylated in either developing or adult tissues from *O. bimaculoides*. This is in stark contrast to vertebrates, where over 80% of CpGs are methylated and to popular invertebrate model organisms, which lack DNA methylation entirely. In octopus, methylated CpGs are clustered in the gene bodies of a subset of transcripts and are absent from transposons and other repetitive sequences. Importantly, this pattern is largely static across developing and adult tissues and correlates with genes that have similar levels of expression across many tissues. In contrast, histone methylation levels are different across tissues in octopus, squid, and bobtail squid. This suggests the intriguing idea that genes which are not highly tissue specific are maintained by DNA methylation whereby those that are dynamically expressed are regulated by a histone code or other epigenetic features.

Evolution, by convergent or stochastic independent events, has selected for variations in the canonical patterns of methylation [3, 12], illustrated by the high level of DNA methylation across the genome of the sponge *A. queenslandica* [17], the clustering of DNA methylation on repetitive elements in the centipede *S. maritima* [88] and the intermediate pattern of methylation in *C. intestinalis*, where it is high on gene bodies and intermediate on repetitive elements [9, 10, 19]. The methylome in *O. bimaculoides* appears be more typical, as it resembles the gene body methylation pattern found in other mollusks [13, 14, 20, 35] and many invertebrates [3, 8, 9, 13, 17]. While gene body methylation is positively correlated with genes expression in organisms ranging from sponges [17, 18] to oysters [13, 20] to mammals [9, 12], it is not clear how this functions. Importantly, we found CpG levels and the pattern of DNA methylation to be nearly identical across adult and developing tissues in *O. bimaculoides* and to be at the same level in all tissues examined in other cephalopods. Interestingly, our finding that the most methylated and the most unmethylated genes in octopus had similar expression patterns across tissues suggests that CpG methylation has the same function in different cell types and argues against this epigenetic modification as a player in the dynamic regulation of gene expression.

In vertebrates, DNA methylation represses transposons [4]. We [89-91] discovered that zebrafish who have lost DNA methylation have multiple severe phenotypes including embryonic lethality, apoptosis, cell cycle defects and innate immune activation [62, 92]. These findings are concurrent with studies in oysters [93] and an annelid worm [18] where blocking DNA methylation causes developmental defects and prevents regeneration. The phenotypes in zebrafish which lack DNA methylation are in part, attributed to activation of TEs [62, 92]. However, while TEs represents about half of the octopus [44], there is a stark contrast between high levels of DNA methylation on TEs in vertebrates and virtual absence of TE methylation in cephalopods and most other invertebrates. This raises the important questions of: what represses TEs in these species and, if DNA methylation is not regulating TEs in octopus, what is it’s function? Answering these questions will require experiments where the DNA methylation machinery can be manipulated in these animals.

In contrast to the consistency of overall level and pattern of DNA methylation across octopus tissues, we found that histone modification levels differ, suggesting a specific histone code regulates the tissue-specific transcriptome. The report of H3K4, H3K9 and H3K36 methylation changes during oyster development [94] suggests that these are dynamic and important regulators of mollusk development. An interesting study profiling H3K4me1/me2/me3, H3K36me3 and H3K27Ac in anemone showed that the epigenetic landscape was similar to that found in bilaterians, with conserved regulation for enhancers in this species [95]. Moreover, the finding that the histone pattern in organisms with diverse DNA methylomes recapitulates the vertebrate pattern – such as the planarian *Schmidtea mediterranea*, which lacks DNA methylator complexes and DNA methylation, as well the highly methylated sponge *A. queenslandica* [96, 97] – suggests that many organisms decouple these different marks in patterning the epigenetic landscape. Future studies integrating histone and DNA methylation profiling with transcriptomics in cephalopod tissues can address how these patterns are integrated.

This work uncovers previously unknown features of the octopus epigenome, which can expand our understanding of epigenetic functionality beyond the few species that are used as a paradigm for knowledge in this field. Studying non-model organisms opens new challenges. For instance, annotation of epigenetic modifiers is based on functions that homologous proteins have in mammals and a thorough functional investigation for each homologous protein should be performed to unequivocally determine their activity in cephalopods. Unfortunately, size and temperate life cycle features of *O. bimaculoides* limits its possibilities for genetic manipulation. Squid, where genetic engineering has been demonstrated [46], and other cephalopod species which are actively being developed as genetic models [47] can serve as alternatives. Identifying the epigenetic patterns of octopus is a first important step in deciphering the how these patterns function to regulate the extraordinary features of these animals will be important future steps.

## Acknowledgments

We gratefully acknowledge the staff in the Marine Resources Center at the Marine Biological Laboratory and in NYUAD Genomics Core Technology Platform, especially Nizar Drou for assistance with annotating the octopus genome. Patrice Delaney and Catherine Palmer provided assistance with the sample collection, Anjana Ramdas Nair edited the manuscript, and all the members of the Sadler lab provided insightful discussion. We especially thank Kirsten Peramba for assistance in octopus care and sample collection and Jan Hsiao and Lola Deng for assistance in building the phylogenomic pipeline. Funding was from the National Science Foundation, NIH R21 (MH119646), HFSP Research Program Grant (RGP0060/2017), Vetlesen Foundation Fellowship (to EE) and the NYUAD Faculty Research Fund (to KCS).

## Author Contributions

The project was conceived and designed by KCS and FM, samples were collected by EE and KCS, analysis was carried out by FM and EE; FM, EE and KCS wrote the paper.

## Supplemental Figure legends

**Supplemental Figure 1. Representative images of *O. bimaculoides***.

**A**. Picture of anesthetized adult male. **B**. Representative image of *O. bimaculoides* arm for collection of distal, medial and proximal samples. **C**. A clutch of 30 dpf hatchlings and **D**. the hatchling used for DNA, RNA and protein extraction.

**Supplemental Figure 2. Distinct biological processes and cellular components of differentially expressed genes octopus tissues**.

GO analysis of **A**. Biological Process and **B**. Cellular Component of each gene cluster identified in Figure 1.

**Supplemental Figure 3. Extended phylogenetic analysis shows high conservation of DNMT1 and UHRF1**.

**A**. Phylogenetic tree of DNMT1 in a representative subset of 50 metazoan and outgroup species. **B**. Phylogenetic tree of UHRF1 in a representative subset of 50 metazoan and outgroup species.

**Supplemental Figure 4. *O. bimaculoides* genome encodes conserved features of DNMT1 and UHRF1 but not UHRF2**.

**A**. Similarity for each domain of *O. bimaculoides* DNMT1 determined by HMMER. Bit score indicates homology score. **B**. Table shows percentage of sequence identity calculated by ClustalOmega for the *O. bimaculoides* DNMT1 protein compared to human, mouse, and zebrafish in a multiple sequence alignment. **C**. Table shows similarity for each domain (with a PFAM ID) of *O. bimaculoides* UHRF1 when compared to the proteome database using HMMER. Bit score indicates homology score. **D**. Table shows percentage of sequence identity calculated by ClustalOmega for the *O. bimaculoides* UHRF1 protein compared to human, mouse, and zebrafish in a multiple sequence alignment. **E**. Similarity for each domain of the *O. bimaculoides* YDG_OCTBM protein determined using HMMER. Bit score indicates homology score. **F**. Percent identity calculated by ClustalOmega for the *O. bimaculoides* YDG_OCTBM protein compared to human, mouse, and zebrafish in a multiple sequence alignment. **G**. Alignment of the SRA domain in *O. bimaculoides* YDG_OCTBM to human, mouse, and zebrafish shows that all the major residues needed for the correct functionality of UHRF1 are not conserved in YDG_OCTBM of *O.bimaculoides*. Alignment to UHRF2 shows no conservation of critical residues between YDG_OCTBM in *O. bimaculoides* and UHRF2 in human, mouse. Residues functionality is based on mouse orthologs

**Supplemental Figure 5. Structural modelling of DNMT1 and UHRF1**.

**A**. 3D structure of the BAH1, BAH2 and CTD domains of DNMT1 in *M. musculus*. **B**. 3D model of BAH1, BAH2 and CTD domains of DNMT1_OCTBM in *O. bimaculoides*. **C**. 3D structure of SRA domain of UHRF1 in *M. musculus*. **D**. 3D model of SRA domain of UHRF1_OCTBM in *O. bimaculoides*.

**Supplemental Figure 6. Pattern of DNA methylation identified by WGBS and RRBS in octopus tissues**.

**A**. Table describing the number and relative percentage of CpGs covered in the *O. bimaculoides* genome by each technique and sample analyzed. CpGs were classified based on the percentage of methylation as methylated (> 80%) or not methylated (< 20%). **B**. Scaled violin plot of CpGs identified by WGBS and RRBS in Supra E and Sub E brain and in one whole-body 30 dpf hatchling. Numbers on the lines indicate the percent of CpGs that are detected as >80% methylated. **C**. Box plot describing the percentage of methylation of CpGs contained in Repetitive Elements (RE) divided by class in the 30 dpf hatchling. Box-and-whisker plots have a center line at the median, lower and upper hinges correspond to first and third quartiles, and whiskers extend from hinges to largest or smallest values no further than 1.5 × IQR (inter-quartile range), while data beyond the end of the whiskers are outlying points that are plotted individually.

**Supplemental Figure 7. Relationship between DNA methylation pattern gene expression, and gene length in octopus**.

**A**. Heatmap of DNA methylation of all full-length transcripts and 2000 bp upstream and downstream as detected in the Sub E brain WGBS data. Clustering and rank order is dictated by Supra E brain samples shown in Figure 4A. **B**. Heatmap of DNA methylation in hatchling full-length transcripts and 2000 bp upstream and downstream. Clustering and rank order is dictated by Supra E brain samples shown in Figure 4A. **C**. Overall DNA methylation of transcripts (TSS to TTS of each transcript) in Sub E brain and hatchling is regressed against transcripts expression (log2(TPM+1)). Transcripts were grouped by percentile of expression values and each dot represents the average value of DNA methylation for each percentile. **D**. Violin plot displays the distribution of transcript length (as log10(width)) in each methylation pattern. p-values were calculated by unpaired non-parametric Kruskal-Wallis test adjusted with Dunn’s multiple comparisons test. **** indicates p-value adjusted < 0.0001.

**Supplemental Figure 8. Supplemental phylogenetic trees**.

**A**. Phylogenetic tree of EZH2, responsible of H3K27me3 deposition, in a representative subset of 19 metazoan and outgroup species. **B**. Phylogenetic tree of KAT2A, mainly responsible of H3K9ac deposition, in a representative subset of 19 metazoan and outgroup species. **C**. Phylogenetic tree of HADC8, mainly responsible of H3K9ac removal, in a representative subset of 19 metazoan and outgroup species.

**Supplemental Figure 9**.

**Histone methyltransferases, acetyltransferases and de-acetylases have a tissue specific expression pattern in octopus A**. Heatmap of the extended panel of histone methyltransferases. **B**. Heatmap of the main histone acetylation factors, acetyltransferase (writers) and de-acetylases (erasers).

## Supplemental Table legends

**Supplemental Table 1**. Statistical metrics of RRBS. **A**. Statistic values of alignment performed in Bismark. **B**. Read coverage statistics per base analyzed by methylKit.

**Supplemental Table 2**. Reference Gene Set (RGS) sources and details of proteins utilized in phylogenomic pipeline.

**Supplemental Table 3**. Genome sources and details of organisms utilized in phylogenomic pipeline.

**Supplemental Table 4**. Trinotate table containing the putative gene names of the transcriptome of *O. bimaculoides*.

**Supplemental Table 5**. Table of Ensembl transcript IDs contained in each Cluster.

**Supplemental Table 6**. Table of Ensembl transcript IDs contained in each heatmap of epigenetic factors. The mapping across different identifiers is also reported for each transcript.

**Supplemental Table 7**. Table of Ensembl transcript IDs contained in each Pattern.

