## Supplemental Figures for "Epigenetic machinery is functionally conserved in cephalopods"

*O. bimaculoides* Adult

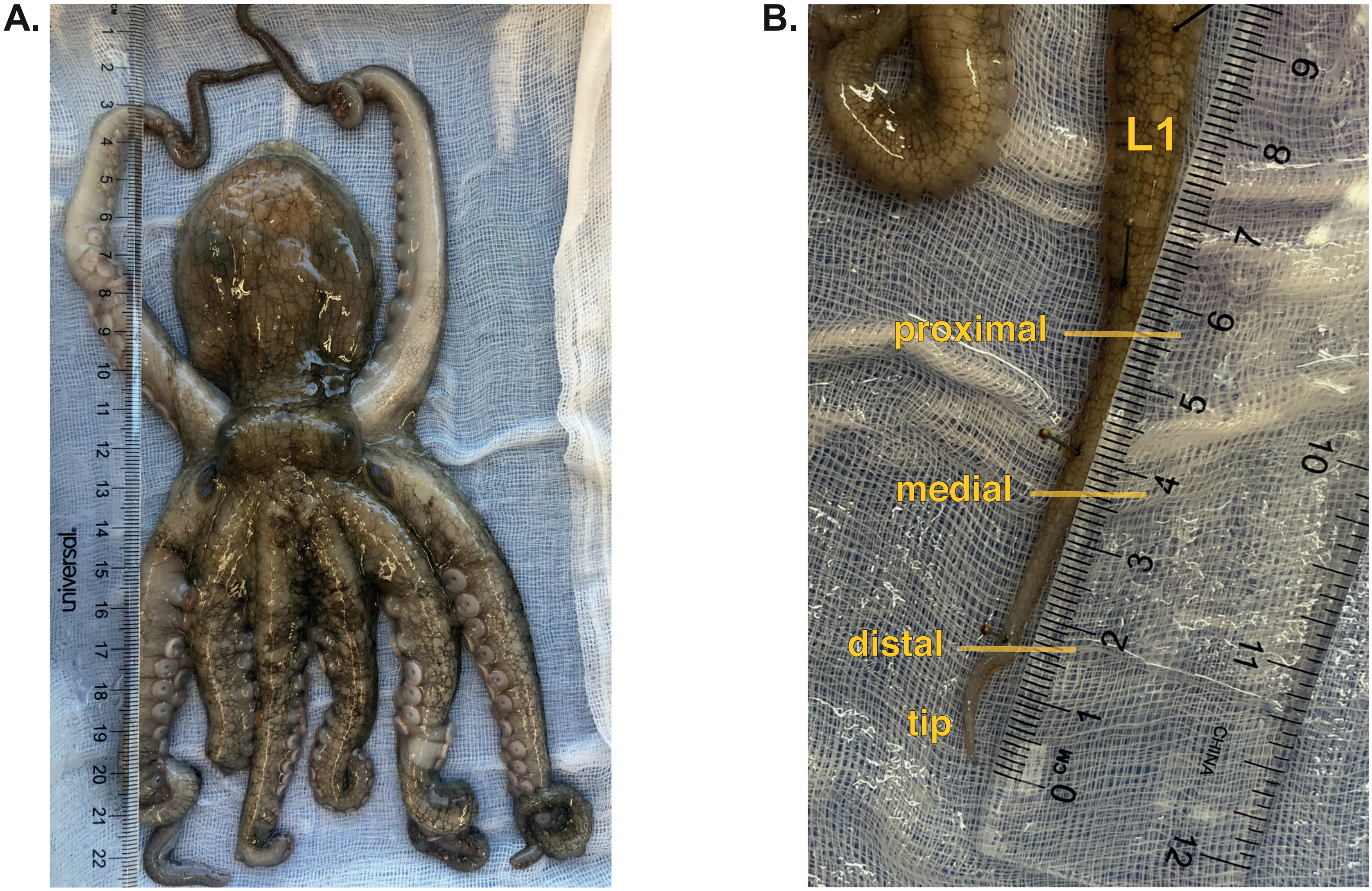


*O. bimaculoides* Hatchling 30 days

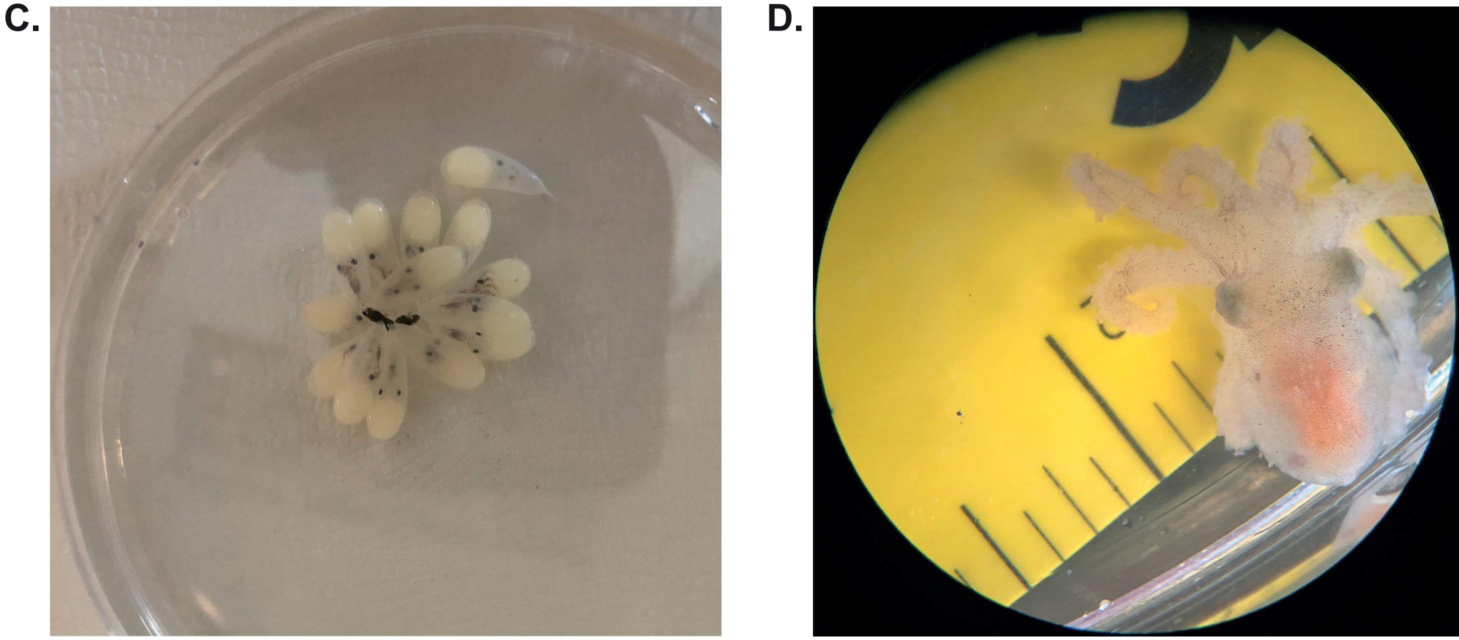


**Suppl. Figure 1**

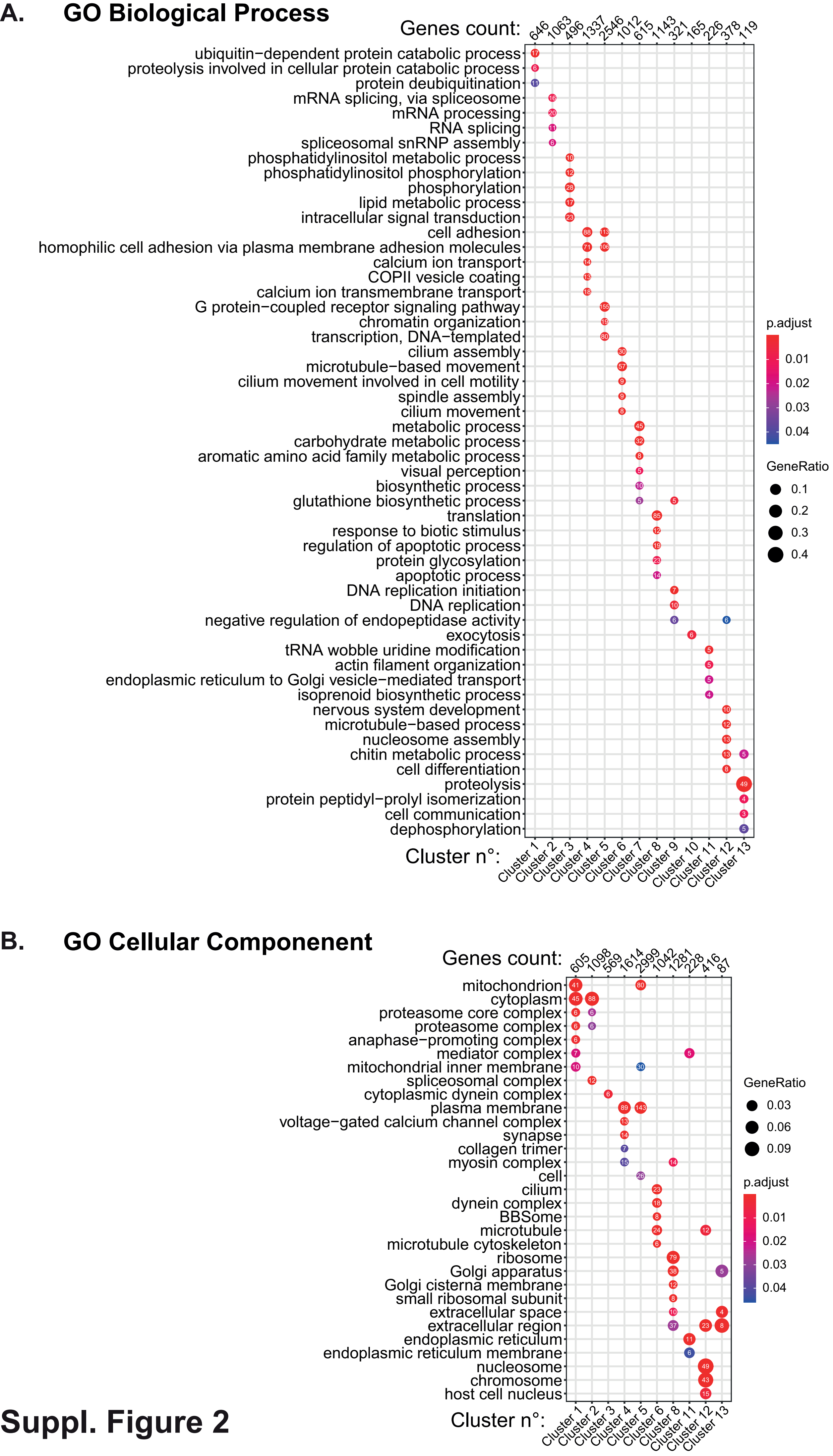


Suppl. Figure 2

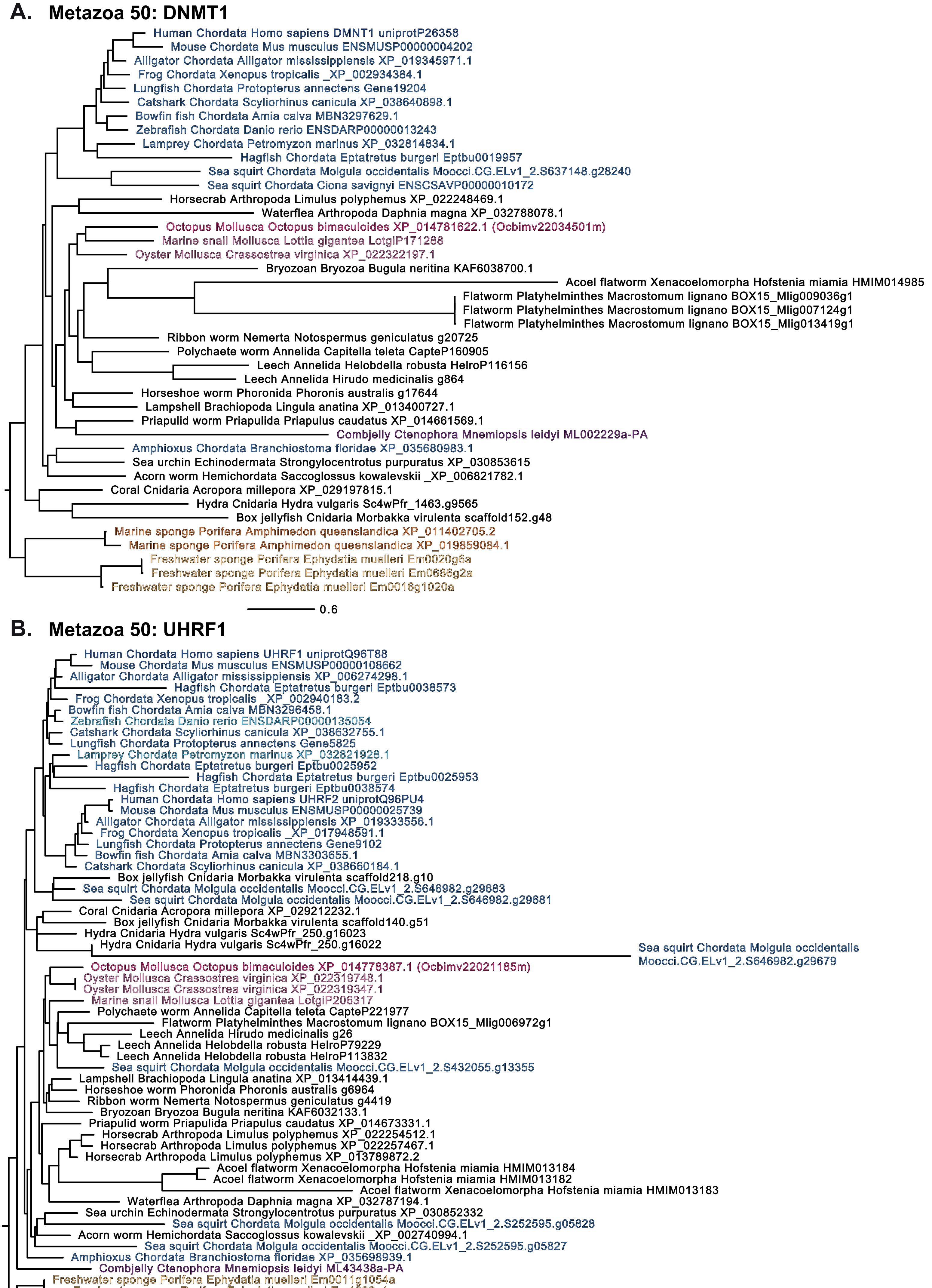


**Suppl. Figure 3**

#### A. DNMT1_OCTBM

| Domain | Pfam ID | Bit Score | Domain E-values | |
|---|---|---|---|---|
| | | | Ind. | Cond. |
| RFTD | DNMT1-RFD | 138.86 | 1.2e-40 | 2.7e-44 |
| CXXC | zf-CXXC | 47.88 | 1.0e-12 | 2.2e-16 |
| BAH1 | BAH | 55.25 | 5.7e-15 | 1.3e-18 |
| BAH2 | BAH | 55.65 | 4.3e-15 | 9.5e-19 |
| CTD (C-5 cytosine methyltransferase) | DNA_methylase | 165.85 | 1.5e-48 | 3.4e-52 |

HMMER (*O. bimaculoides* protein sequence vs. entire protein sequence database)

#### B. DNMT1_OCTBM

| Organism | UniProt ID | Gene ID | % of Identity |
|---|---|---|---|
| Octopus bimaculoides | A0A0L8GEZ1 | *Ocbimv22034501m.g* | 100.00 |
| Danio rerio | Q8QGB8 | *dnmt1* | 56.01 |
| Homo sapiens | P26358 | *DNMT1* | 54.79 |
| Mus musculus | P13864 | *Dnmt1* | 52.24 |

ClustalOmega multiple protein sequence alignment

#### C. UHRF1_OCTBM

| Domain | Pfam ID | Bit Score | Domain E-values | |
|---|---|---|---|---|
| | | | Ind. | Cond. |
| UBL | ubiquitin | 52.61 | 2.8e-14 | 6.2e-18 |
| TTD | TTD | 171.29 | 1.3e-50 | 2.9e-54 |
| PHD | PHD | 47.58 | 1.1e-12 | 2.5e-16 |
| SRA | SAD_SRA | 190.13 | 1.9e-56 | 4.2e-60 |
| RING | Not unique ID | Manually annotated | % identity (count) = 64.1% (39) | |

HMMER (*O. bimaculoides* protein sequence vs. entire protein sequence database)

#### D. UHRF1_OCTBM

| Organism | UniProt ID | Gene ID | % of Identity |
|---|---|---|---|
| Octopus bimaculoides | A0A0L8IC59 | *Ocbimv22021185m.g* | 100.00 |
| Danio rerio | E7EZF3 | *uhrf1* | 56.37 |
| Homo sapiens | Q96T88 | *UHRF1* | 55.88 |
| Mus musculus | Q8VDF2 | *Uhrf1* | 53.55 |
| Homo sapiens | Q96PU4 | *UHRF2* | 50.19 |
| Mus musculus | Q7TMI3 | *Uhrf2* | 50.00 |

ClustalOmega multiple protein sequence alignment

#### E. YDG_OCTBM

| Domain | Pfam ID | Bit Score | Domain E-values | |
|---|---|---|---|---|
| | | | Ind. | Cond. |
| SRA | SAD_SRA | 195.46 | 4.3e-58 | 2.4e-62 |

HMMER (*O. bimaculoides* protein sequence vs. entire protein sequence database)

#### F. YDG_OCTBM

| Organism | UniProt ID | Gene ID | % of Identity |
|---|---|---|---|
| Octopus bimaculoides | A0A0L8H8G0 | *Ocbimv22020196m.g* | 100.00 |
| Mus musculus | Q8VDF2 | *Uhrf1* | 34.46 |
| Homo sapiens | Q96T88 | *UHRF1* | 33.53 |
| Danio rerio | E7EZF3 | *uhrf1* | 32.42 |
| Mus musculus | Q7TMI3 | *Uhrf2* | 31.69 |
| Homo sapiens | Q96PU4 | *UHRF2* | 31.08 |

ClustalOmega multiple protein sequence alignment

## G.

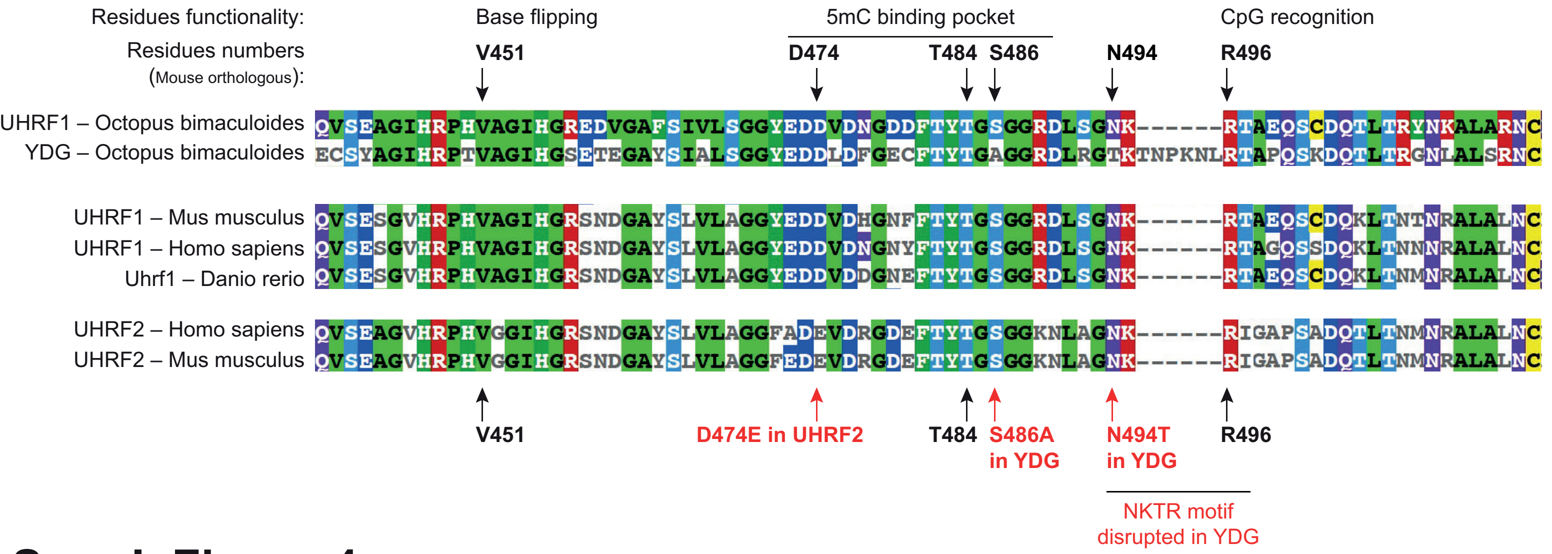


### Suppl. Figure 4

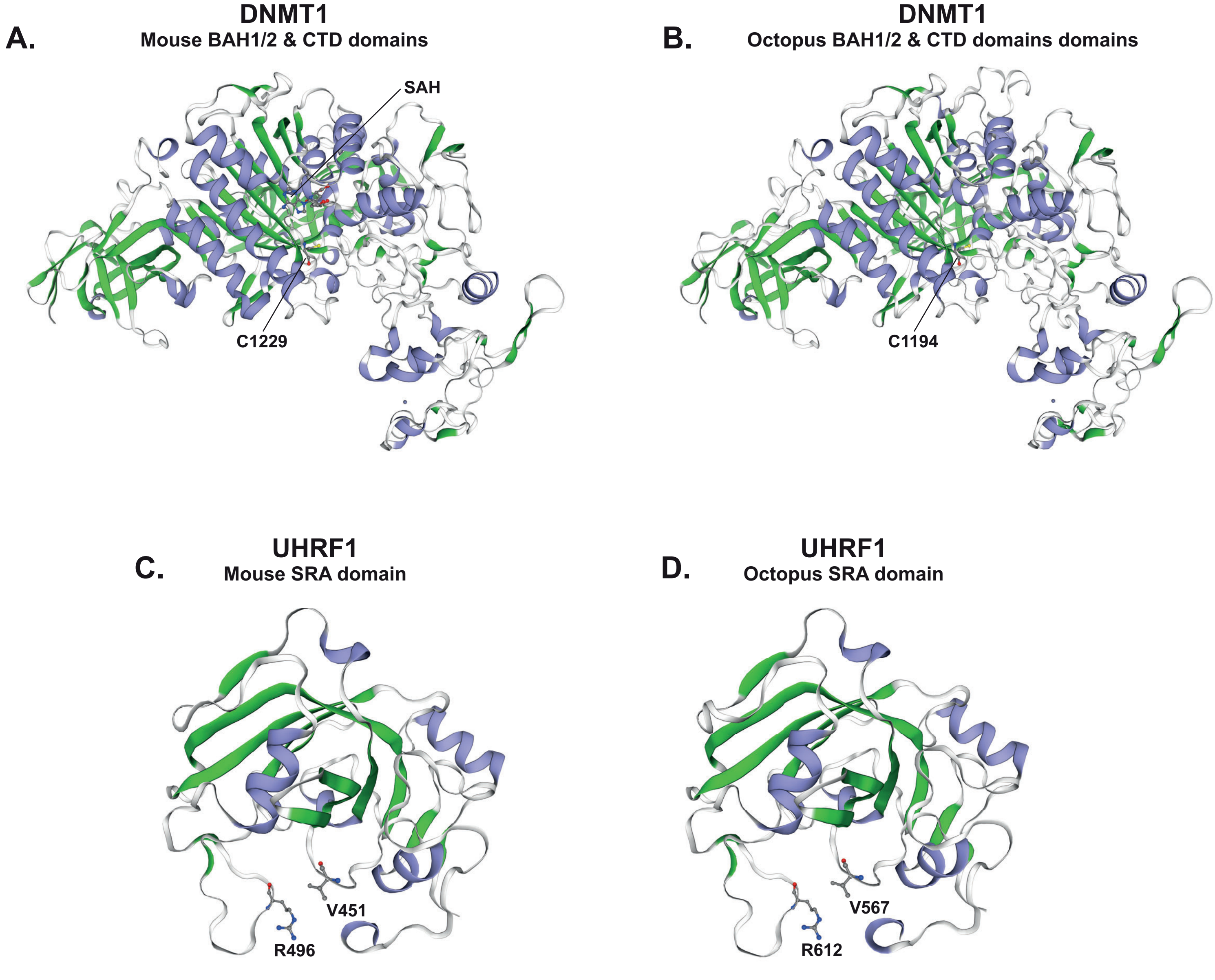


### Suppl. Figure 5

**A.**

| Total CpGs | 80,660,576 | | |
|---|---|---|---|
| **Sample** | **Covered CpGs** | **Methylated CpGs (>80%)** | **Unmethylated CpGs (<20%)** |
| **RRBS Hatchling** | 2,403,266 | 87,223 | 2,172,200 |
| Percent of total | 2.98% | 3.63% | 90.39% |
| **WGBS Supra E** | 64,845,686 | 3,897,277 | 55,918,461 |
| Percent of total | 80.39% | 6.01% | 86.23% |
| **WGBS Sub E** | 47,053,504 | 3,542,069 | 40,626,272 |
| Percent of total | 58.34% | 7.53% | 86.34% |

**B.**

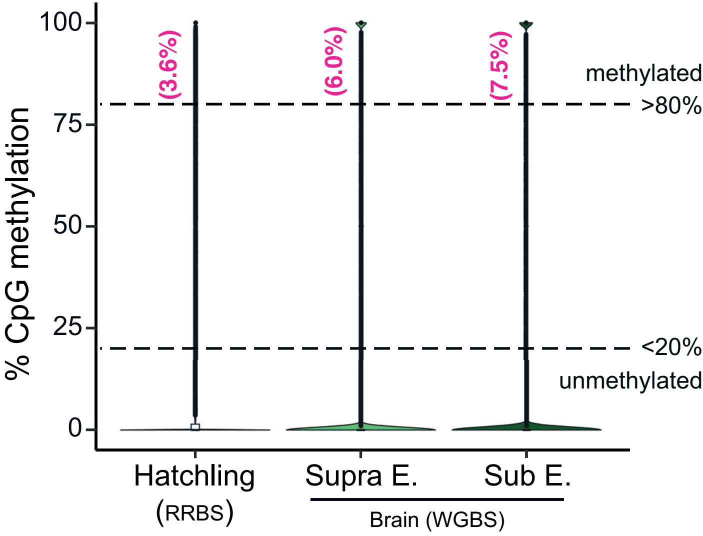


**C.**

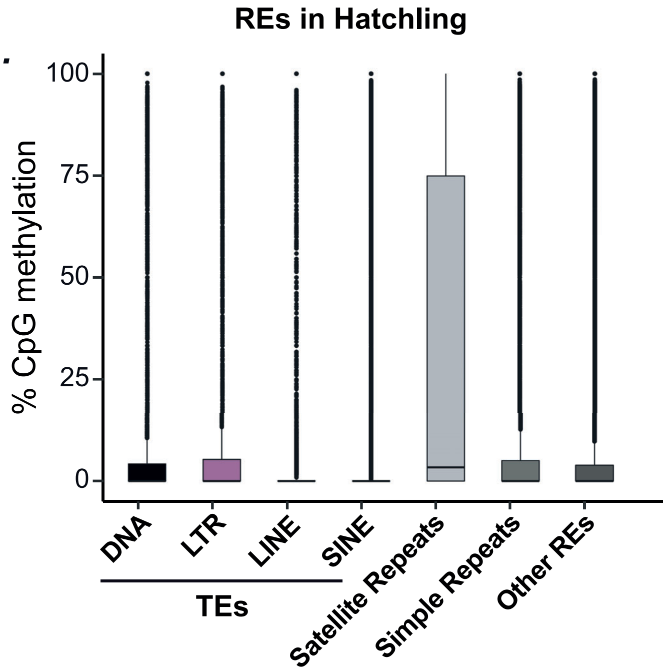


**Suppl. Figure 6**

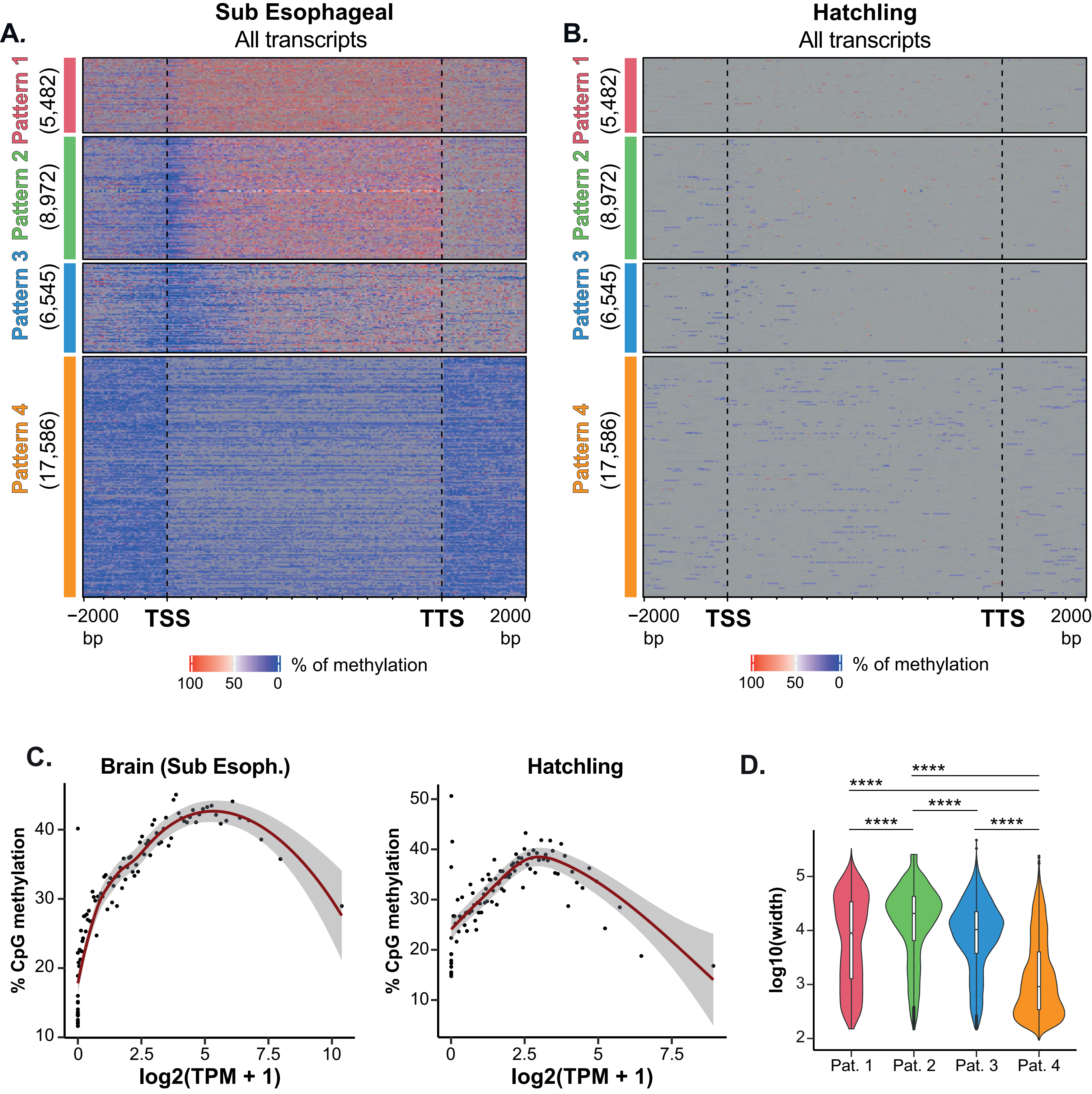


Suppl. Figure 7

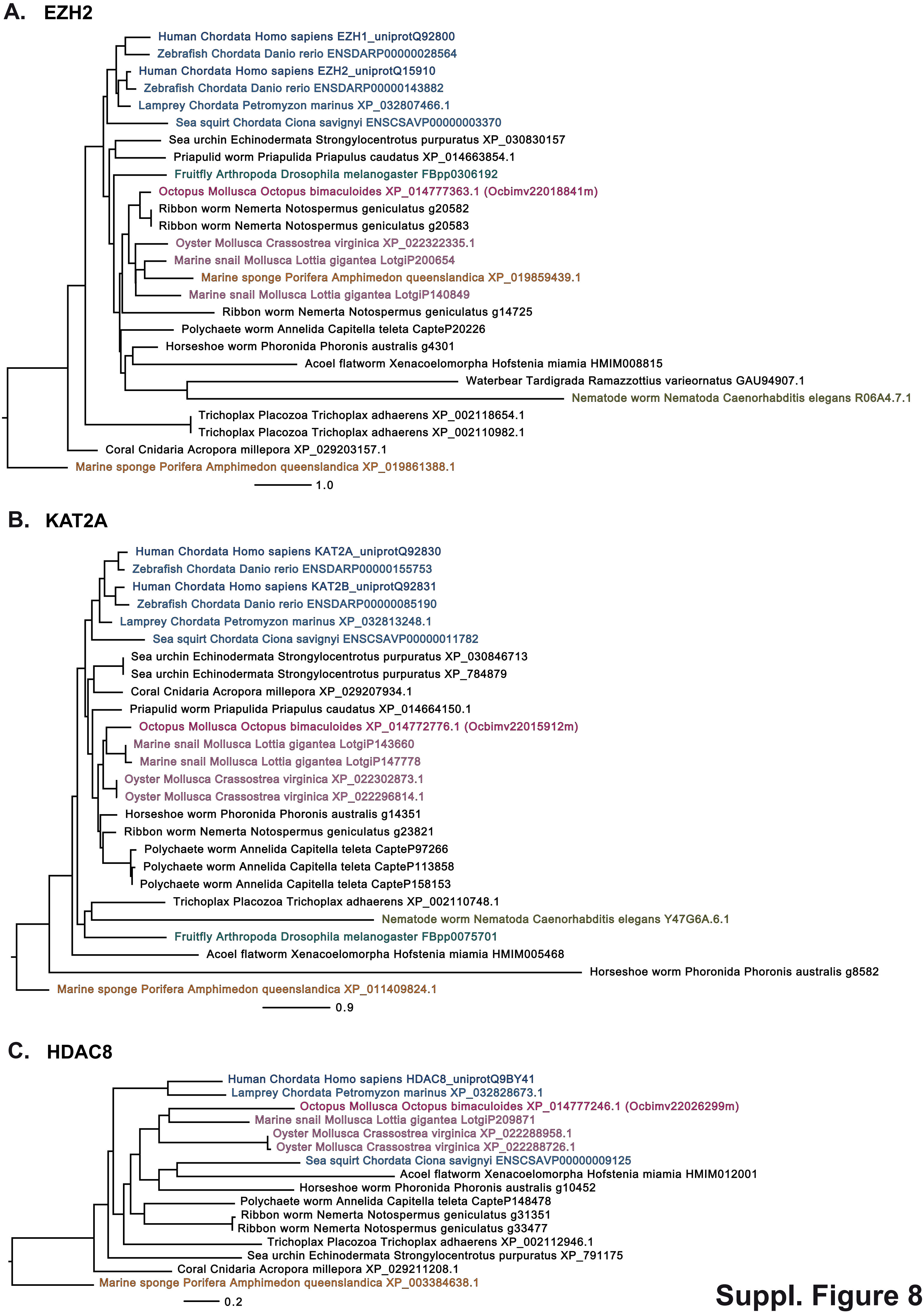


**Suppl. Figure 8**

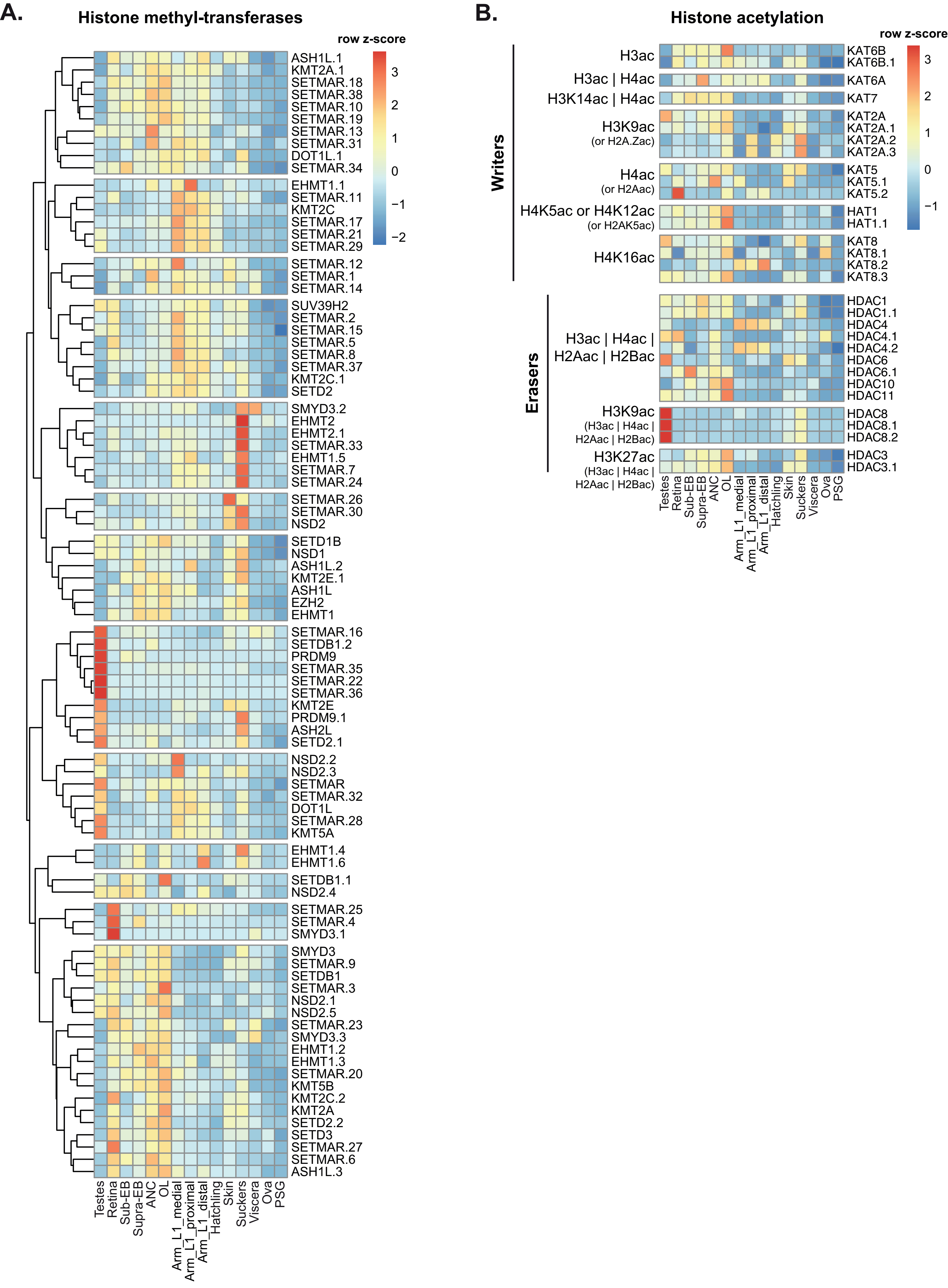


**Suppl. Figure 9**
